# Sudden shifts in expression after small-scale duplication in vertebrates and strong support for the ortholog conjecture

**DOI:** 10.1101/2023.12.29.571877

**Authors:** Tina Begum, Pablo Duchen, Christabel Floi Bucao, Marc Robinson-Rechavi

## Abstract

Gene duplication is a potential source of innovation, but the evolutionary dynamics of functional change are still poorly understood. Under the debated “ortholog conjecture”, most functional change and innovation is assumed to follow duplication. Testing the ortholog conjecture allows to better understand and characterize the way in which gene function evolves. Most models of functional evolution assume continuous change, an assumption which we challenge here. We have applied a Lévy model of evolutionary trait jumps to the evolution of gene expression in vertebrates, with a special focus on duplication in teleost fishes. We show for the first time that trait jumps strongly affects paralogs, in addition to other modes of functional evolution. We find that at least 25% of teleost fish small-scale duplicates follow a rapid evolutionary rate shift model for both expression level and tissue-specificity, much more than after speciations. However, genome-wide duplicates (ohnologs) do not support such a trait jump model, and thus follow a different evolutionary dynamic. While there is some evidence for more positive selection at the protein-coding level after duplication, it is not strongly linked to jumps in expression. Finally, both small-scale paralogs and ohnologs strongly support the ortholog conjecture by contrasting speciation branches pre- and post-duplication to the duplication branches themselves, with trait jumps explaining much of the higher phylogenetic independent contrasts between small-scale paralogs.

**Significance statement:** The debate on the ortholog conjecture, i.e. that gene function changes little between orthologs but changes frequently between paralogs, provides a framework to understand better the evolution of gene function. Here we add two pieces to the puzzle: a novel way to use phylogenetic contrasts to test the ortholog conjecture, by comparing not only duplication to speciation, but speciation according to whether they were preceded by a duplication; and a model of jumps rather than continuous change of gene function. We tested these on vertebrates, with emphasis on teleost fishes, distinguishing small-scale duplications and whole-genome duplication; in all cases we support strongly the ortholog conjecture. We find that trait jumps strongly affect small-scale paralogs but not genome duplication paralogs, providing an exciting new model for gene function evolution.

## Introduction

Gene duplication is a potential source of functional innovation, and plays an important role in the evolution of genomes (Ohno 1970; Chen and Zhang 2012; Guschanski et al. 2017). Such innovation can lead to overall function gain, e.g. through neofunctionalization or subfunctionalization, or to specialization, and can be symmetric or not between paralogs (Assis and Bachtrog 2013; Assis and Bachtrog 2015; Braasch et al. 2016; Lien et al. 2016; Guschanski et al. 2017; Sandve et al. 2018). Paralogs can also be retained without any functional change, due to dosage constraints (Gout and Lynch 2015; Thompson et al. 2016). In this context, there has been a debate about the long standing hypothesis that paralogs diverge generally more in function than orthologs, the “ortholog conjecture” (Koonin 2005; Studer and Robinson-Rechavi 2009; Nehrt et al. 2011; Altenhoff et al. 2012; Chen and Zhang 2012; Gabaldón and Koonin 2013; Rogozin et al. 2014; Kryuchkova-Mostacci and Robinson-Rechavi 2016; Dunn et al. 2018; Fukushima and Pollock 2020; Stamboulian et al. 2020; Begum and Robinson-Rechavi 2021). This ortholog conjecture is of interest both in practice, for functional annotation of homologs, and in principle, to understand how gene function evolves. Thus contrasting the evolution of genes after duplication or speciation provides a clear framework to investigate the evolution of function.

Since the retention rates and evolutionary dynamics of duplicates vary among lineages and between genes, they need to be studied using a phylogenetic framework with proper care (Pagel 1999; Blomberg et al. 2003; Dunn et al. 2018; Begum and Robinson-Rechavi 2021). A phylogenetic approach also provides the opportunity to explicitly test different models of evolution. In most studies, the implicit model is that function evolves gradually, with changes in evolutionary rate mapped to specific branches of the phylogeny. These approaches, while important, remain less explored than simpler pairwise comparisons.

There is abundant evidence for an asymmetric steady acceleration of sequence-level evolutionary rates (e.g. *d*N/*d*S) after duplication (Conant and Wagner 2003; Gu et al. 2005; Brunet et al. 2006; Kim and Yi 2006; Cusack and Wolfe 2007; Scannell and Wolfe 2008; Studer and Robinson-Rechavi 2009; Panchin et al. 2010; Pegueroles et al. 2013; Roselló and Kondrashov 2014; Braasch et al. 2016; Lien et al. 2016; Holland et al. 2017; Sandve et al. 2018). However, functional evolution following gene duplication might not be limited to such gradual modes of evolution; it might not even be the main mode of divergence. Studies of phenotype evolution have found that there can be a period of instantaneous bursts or jumps in trait values at the bases of clades that transitioned into new adaptive zones, due to the appearance of an evolutionary novelty (Simpson 1944; Bokma 2008; Landis et al. 2013; Duchen et al. 2017; Landis and Schraiber 2017). This pattern of evolutionary innovations is called “pulsed evolution”, where the background trait evolutionary rates σ^2^ remain unaltered following a jump with a strength λ (Duchen et al. 2017; Gao and Wu 2022). Such non-gradual evolutionary jumps were identified with a frequency of once-per-million-years in vertebrate evolution, using a pure-jump model (Uyeda et al. 2011; Landis et al. 2013). It is still an open question whether such pulsed evolution exists for gene and genome duplicates (Begum et al. 2021). We propose here to examine this question in light of the ortholog conjecture.

Since phenotypic evolution may arise through mutations that affect gene expression patterns (Guschanski et al. 2017), and that these are readily comparable between genes and species, we have performed large-scale phylogenetic comparative analyses considering two phenotypic traits which characterize gene expression: average expression levels, and tissue specificity (τ). We focused on teleost fishes since they have undergone a whole genome duplication (the “FishWGD”) around 320 million years ago (Mya), followed by a spectacular evolutionary radiation leading to more than half of all vertebrate species (Pasquier et al. 2017; Davesne et al. 2021; Parey et al. 2022). This allowed us to address the following questions using a phylogenetic framework: Do we find a global support for the ortholog conjecture for different types of duplicates and different expression traits? What is the contribution of instantaneous trait jump mechanisms to paralog evolution? We also explored whether changes in expression of paralogs are linked to positive selection at the coding-sequence level.

## Results

### Phylogenetic independent contrasts support the ortholog conjecture robustly

Phylogenetic Independent Contrasts is a phylogenetic comparative method which is well suited to test whether gene duplication promotes more functional divergence than evolution of genes in the absence of duplication, if care is taken in its application (Dunn et al. 2018; Begum and Robinson-Rechavi 2021). We used gene expression as a proxy for function in vertebrates (Fig. 1), through two quantitative traits: i) tissue specificity, τ (Yanai et al. 2005), and ii) average expression level. Gene expression is a common proxy for gene function because it is amenable to measurement over all genes in several species, and the level and tissue-specificity of expression patterns correlates well with biological function of genes (Kryuchkova-Mostacci and Robinson-Rechavi 2017). To compute average gene expression levels, we normalized gene expression levels such that the cross-species expression data clearly separates into clusters per organ (Supplementary figs. S1A and S1B)). This way gene expression is comparable across species and tissues or organs (Brawand et al. 2011; Fukushima and Pollock 2020).

**Figure 1:**
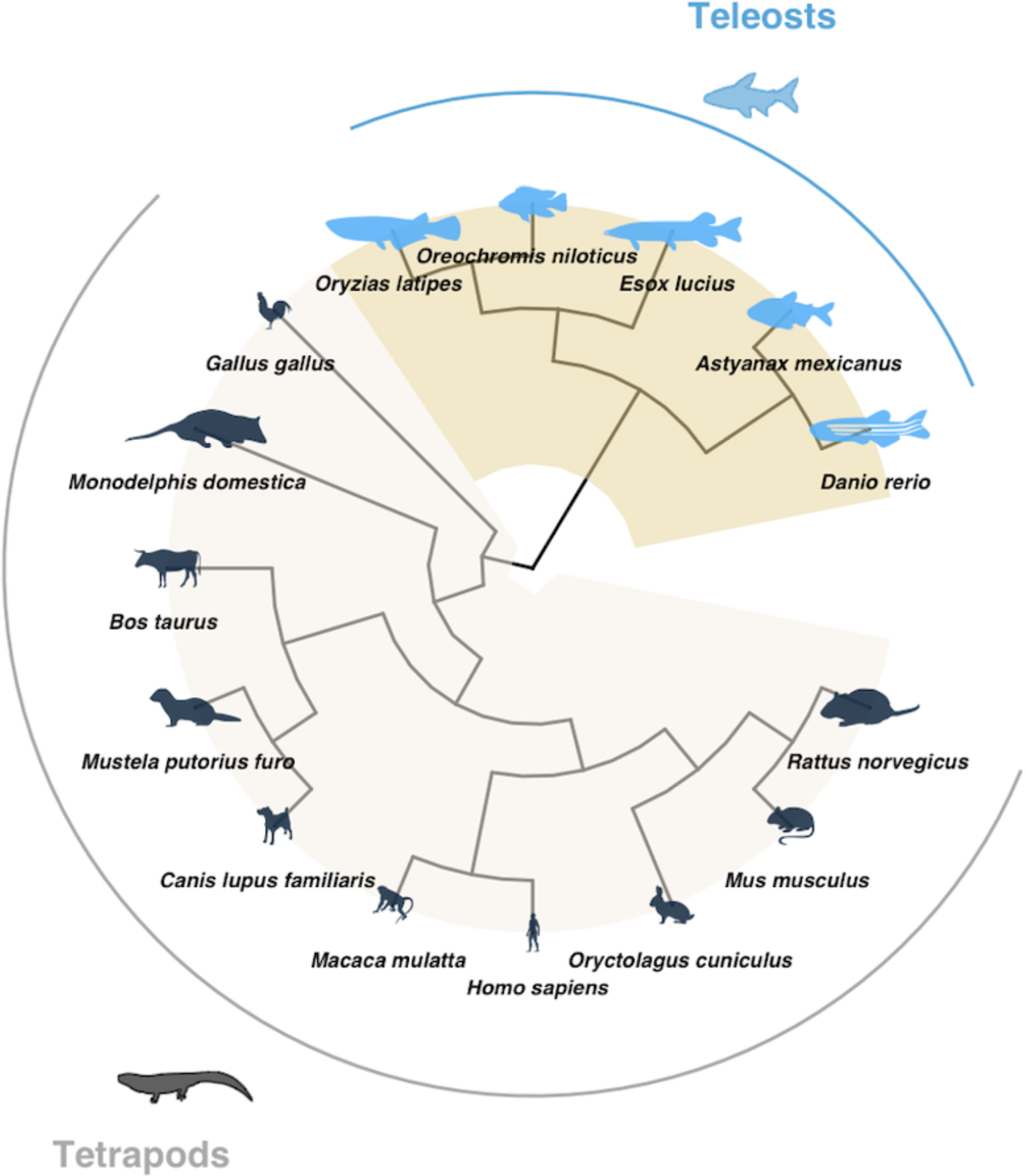
The vertebrate species phylogeny of this study. The ‘brownish’ color block highlights 5 teleost fishes, and the ‘off-white’ color block highlights the 10 tetrapods.

The phylogenetic independent contrast of each node in a phylogenetic tree reflects independent changes in continuous traits, where higher contrasts capture more evolutionary changes in traits (Felsenstein 1985; Dunn et al. 2018; Begum and Robinson-Rechavi 2021; Begum et al. 2021). If duplication favors trait diversification, we expect higher Phylogenetic Independent Contrast values (PICs) for duplications than for speciations. This is indeed the case for both gene expression traits, and both for standardized phylogenies restricted to teleosts or including all 15 vertebrates of this study (Supplementary fig. S2). These results support the ortholog conjecture for vertebrates. In our analyses, since each data point is a node of a tree, it is possible that gene families with many duplication nodes bias the ortholog conjecture test (Begum and Robinson-Rechavi 2021). Hence, the difference observed could be solely due to a small number of gene families with large numbers of duplications. To verify, we first computed the mean PIC per event for each tree to have one datapoint per gene family, along with the frequency of each event. We then restricted our analysis to gene trees with a maximum of five duplication events, with no limitation in speciation events (Supplementary fig. S3A). This accounts for more than 94% of total gene trees for both the traits. Our support for the ortholog conjecture still holds (Supplementary fig. S3B). This suggests that our observation was not solely due to a small number of gene families with large numbers of duplications.

Further analyses on synteny validated teleost third-round whole genome duplicates (FishWGDs) or strict ohnologs, and on strict small-scale duplicates (SSDs) in teleosts (Figs. 2A-2D) also support the ortholog conjecture for both gene expression traits. So do similar analyses on all 15 vertebrates (Supplementary figs. S4A-S4D). We observed a much weaker impact of genome duplications than of small-scale duplications on gene expression trait divergence in terms of effect size. This could be because the ohnologs are much older than the small-scale paralogs (median age of ohnologs: 332 Mya, median age of SSDs: 103 Mya, Wilcoxon two-tailed tests *P* < 2.2 × 10^-16^), or because of differences in evolutionary dynamics.

**Figure 2:**
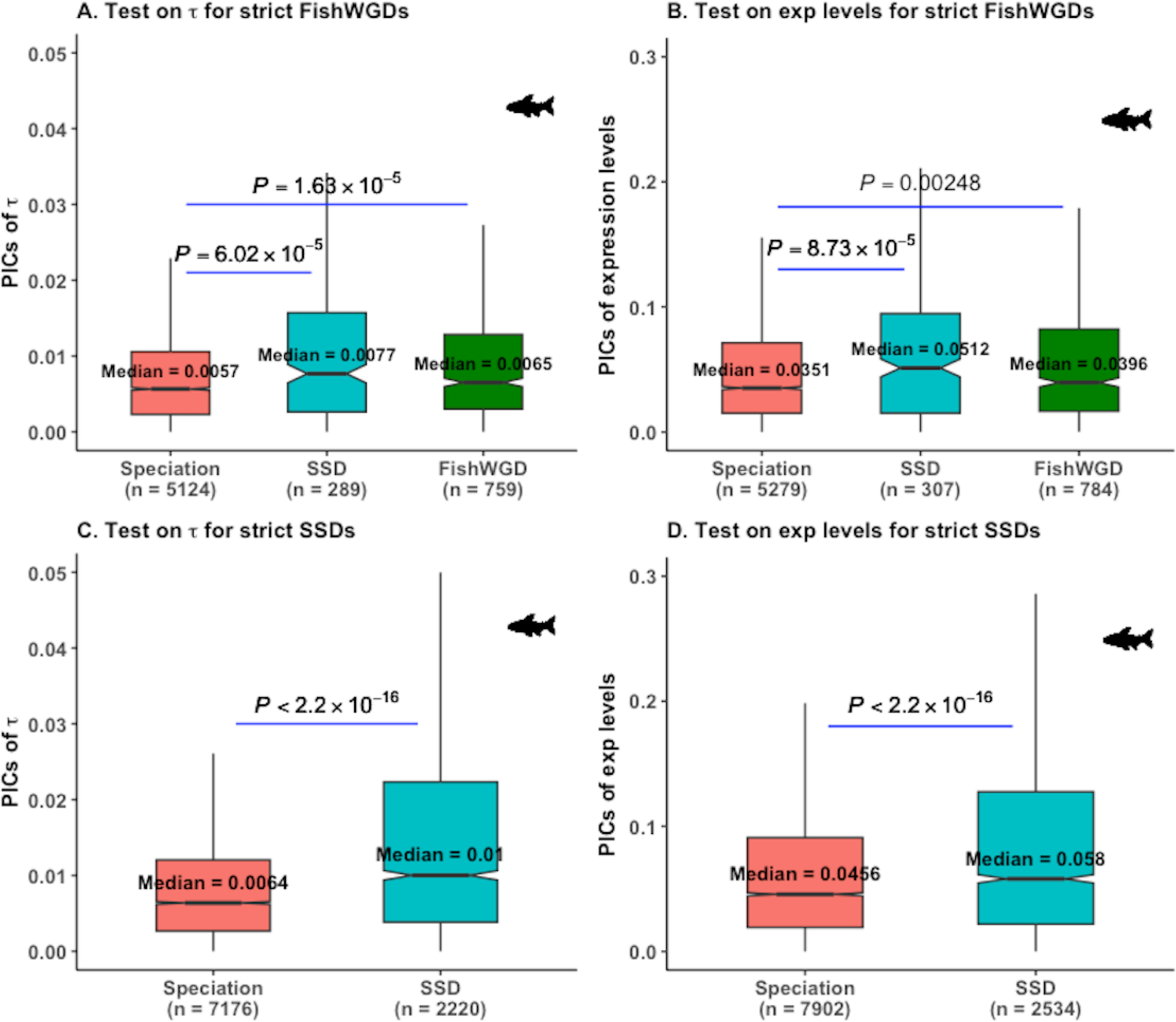
Tests of the ortholog conjecture by phylogenetic independent contrasts according to type of duplication in teleosts. 746 and 774 trees with whole genome duplications were used in (A)-(B), while 2482 and 2717 trees with small-scale gene duplications were used in (C)-(D). PICs: Phylogenetic independent contrasts, FishWGD(s): Fish specific 3R whole genome duplicate(s) (or ohnologs), SSD(s): small-scale duplicate(s). PICs of teleosts specific nodes were used for these analyses. *P* values are from Wilcoxon two-tailed tests. Since trees with whole genome duplicates may also contain SSDs apart from strict FishWGDs, both types of duplicates are used for analyses in (A)-(B).

If the effect of duplications is time-dependent (Gu et al. 2005; Begum et al. 2021), evolution along speciation branches following duplication branches might be impacted by the duplication. This could impact testing of the ortholog conjecture if we simply compare all duplication branches to all speciation branches. To account for this effect, we compared contrasts not only between speciation and duplication nodes, but also between speciation nodes prior to duplications (pre-duplication speciations or “ancestral” speciations) and speciation nodes immediately following duplications (post-duplication speciations or “descendant” speciations) (Fig. 3A). Pre-duplication speciation branches are devoid of any effect of duplication, and thus comparisons of pre-duplication speciation contrasts to its subsequent duplication contrasts provide a more specific test of the ortholog conjecture. Moreover, comparisons of pre- and post-duplication speciation branches provide insight into the long term effect of duplications. We first restricted this comparison to the oldest duplication node along a path in each tree.

**Figure 3:**
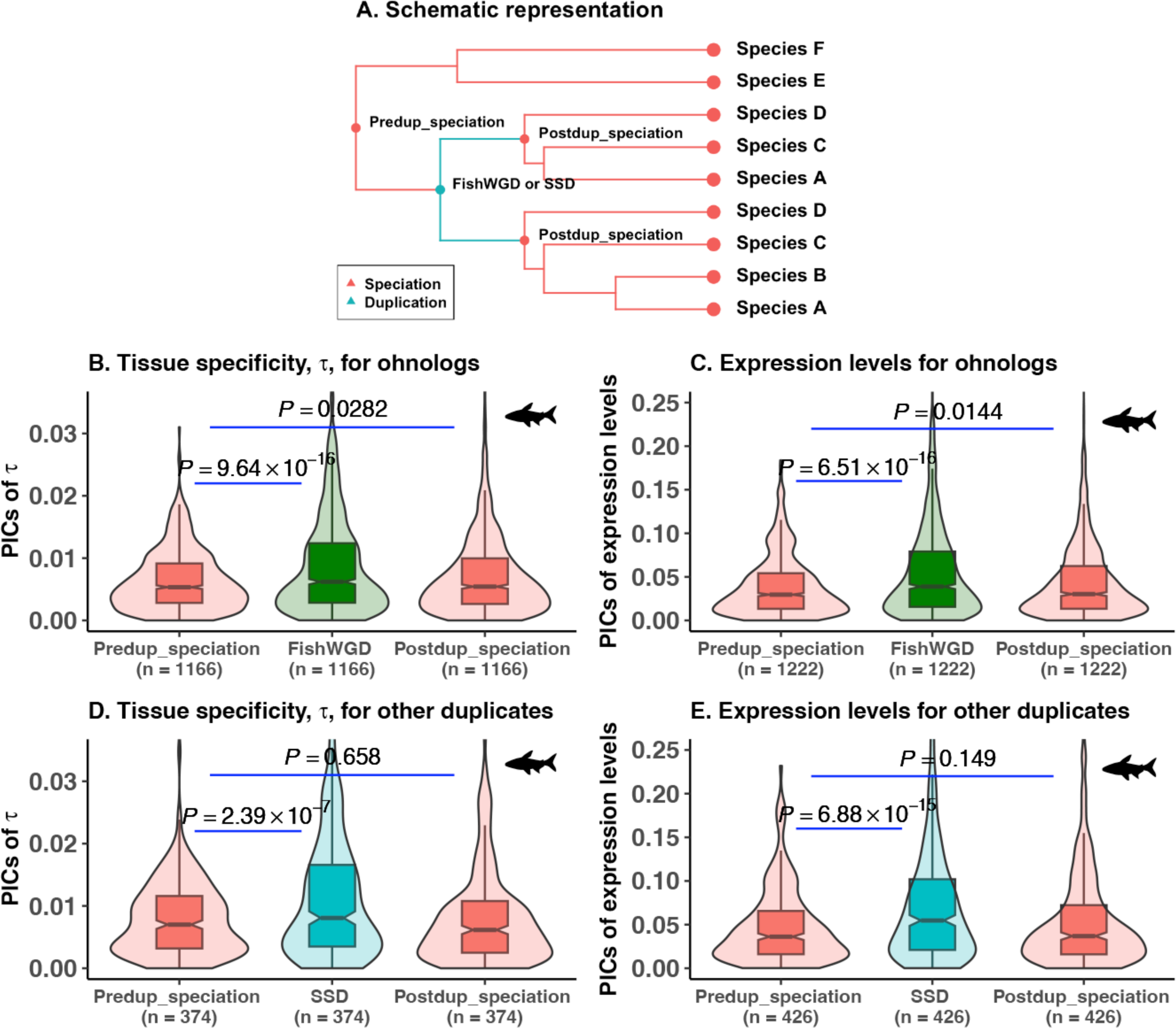
Tests of the ortholog conjecture by phylogenetic independent contrasts including speciations before and after duplications in teleosts. (A) Illustration of how we can test the impact of a duplication by comparing duplication nodes with speciation nodes before and after the duplication. (B)-(E) 581, 610, 339 and 377 trees passing our criteria of Fig. 3A, respectively were used for the analyses. We used the oldest duplication node along a path of a phylogeny for identification of pre-duplication and post-duplication speciations. We considered the contrasts of teleosts specific gene or genome duplication and descendant speciation nodes in (B)-(E) in absence of teleost-specific pre-duplication speciation nodes contrasts. In many cases, we obtained two or three post duplication speciation events following a single duplication event. Therefore, the number of nodes used for comparisons in the plot increased relative to the number of gene trees. PICs: Phylogenetic independent contrasts, exp: expression, Predup_speciation: pre-duplication speciation, Postdup_speciation: post-duplication speciation, FishWGD: fish specific 3R whole genome duplication, and SSD: small-scale duplication. Since contrasts are estimated for each tree, paired Wilcoxon tests were used to compare them.

The contrasts of duplications were always significantly higher than those of ancestral or descendant speciations (Figs. 3B-3E, Supplementary fig. S5). In addition, there was a trend for descendant speciations to have higher contrasts than ancestral speciations, indicating a longer-term impact of the duplication on evolutionary rates. Depending on duplication type and species sampling, this difference was sometimes but not always significant. This might be due in part to small effect sizes which are difficult to test on small subsets of the data, e.g. only ∼14% of the 4139 strict SSD trees fit the criteria of Fig. 3A; and in part to large divergence times between the FishWGD event and the next speciations. To assess bias in the empirical data sets, we performed randomization of traits (Begum and Robinson-Rechavi 2021). Randomized traits showed a pattern distinct from that of real data in all cases (Supplementary figs. S6 and S7), indicating a biological pattern rather than the sole result of a bias in the data (Begum and Robinson-Rechavi 2021). Indeed after randomization the contrasts no longer follow a Brownian model, and become dependent on phylogenetic parameters such as node depth or node age. Hence, the contrasts increase as the node depths decrease towards present time along the randomized phylogenies (Supplementary figs. S6 and S7). Contrasts in randomized phylogenies increase with decreasing node depth, from pre-duplication speciation to duplication to post-duplication speciation nodes, showing that bias is a real concern. Yet unlike in real data, on randomized trees duplication PICs are not higher than post-duplication speciation PICs, showing that the higher PIC of duplications is not due to such biases.

In case of the youngest duplication node along a path, time-dependent effects of prior duplication(s) might impact the contrasts of pre-duplication speciation branches of the youngest duplication branches. Our result was robust when we considered the youngest duplication node along a path whenever available (Supplementary figs. S8 and S9). This indicates that the pattern we observed was not strongly affected by the timing of duplication nor by the risk of a few wrongly annotated genome duplication nodes. Overall, we find that phylogenetic contrasts support the ortholog conjecture in vertebrate gene expression evolution.

### Lévy’s trait jump: sudden shifts in gene expression after small-scale duplication

Continuous traits can change in evolution in sudden shifts or “jumps” in values, without a change of the evolutionary rate outside of this jump. Although Gaussian processes (*i.e.*, Brownian motion (BM) and Ornstein–Uhlenbeck (OU) process) have been widely used to model continuous trait evolution in duplicates (Fukushima and Pollock 2020; Begum and Robinson-Rechavi 2021; Gillard et al. 2021), the relevance of rapid trait jumps to gene and genome duplicates has not been studied. If such jumps were common in gene evolution, this would impact characterization of the ortholog conjecture. Hence, we applied the method “levolution” (Duchen et al. 2017; Duchen et al. 2021) that couples a Brownian model with occasional jumps occurring as a Poisson process, to infer the locations and strength of evolutionary jumps for gene expression traits (Fig. 4A and 4B). Under the ortholog conjecture, our expectation is that there should be more jumps in traits after duplications than after speciations.

**Figure 4:**
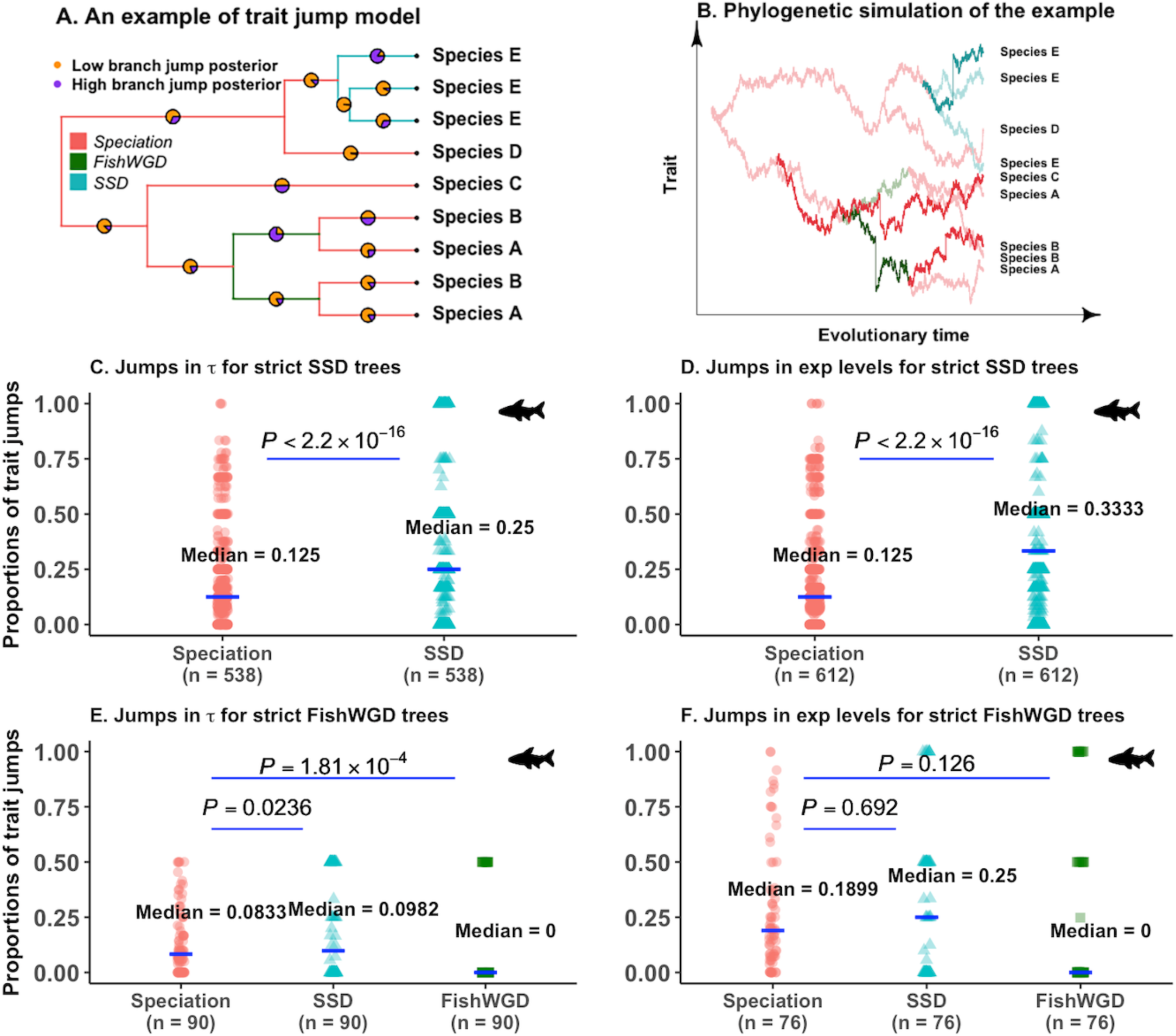
Trait jump models for tissues-specificity and for average gene expression levels contrasting types of duplication in teleosts. FishWGD: fish specific 3R whole genome duplication, SSD: small-scale duplication, and exp: average gene expression levels. (A) An example of a FishWGD tree to demonstrate the branch-specific posterior probability of trait jumps as inferred by levolution (Duchen et al. 2017; Duchen et al. 2021) in a phylogenetic tree. (B) Phylogenetic simulation following the example tree of Fig. 4A. Colors show branches belonging to different events as shown in Fig. 4A. Branches following jumps in their traits are shown by darker shades. (C)-(E) Proportion of trait jumps according to type of duplications and to expression traits. We used a strict branch jump posterior cutoff of ≥ 0.7 to consider a rapid shift in trait values. Only teleosts-specific clades were used for the analyses. We used 1557 and 1593 strict SSD trees in (B) and (C), and 435 and 398 strict FishWGD trees in (D) and (E), respectively showing at least one branch jump with the above-mentioned threshold along a phylogeny for our analyses. For the comparison of the proportions of trait jumps, we used trees with at least one teleost-specific speciation and duplication. Since proportion of shifts in traits per branch of events is estimated for each tree, paired Wilcoxon test is used to compare the difference.

First, we do detect jumps in the evolution of gene expression, affecting more than half of all gene trees (Figs. S10A and S10B), using a branch jump posterior threshold of 0.7. We used a threshold of 0.7 to detect jumps with sufficient accuracy, because uncertainty increases when the posterior probability value decreases (Lemmon and Moriarty 2004; Revell 2014; Pilmann Kotěrová et al. 2024). Second, there are indeed more than twice as many jumps on duplication branches than on speciation branches for small scale duplications in teleosts (Figs. 4C and 4D). Third, there are almost no jumps after whole genome duplication in teleosts (Figs. 4E and 4F). There are jumps on the speciation and SSD branches of WGD trees, thus the lack of jumps on WGD branches is not due to a bias in WGD genes. If different types of duplications are not distinguished, in a naive testing of the ortholog conjecture, there is overall a significantly higher proportion of jumps per branch for both the gene expression traits in teleosts and in vertebrates following duplications than speciations (Supplementary figs. S10C-S10F). Such rapid trait shifts are slightly more asymmetric for τ (asymmetric *vs.* symmetric jump branches: 51.44% *vs.* 48.56%, two-sample test for equality of proportions, χ^2^test, *P* = 4.05 × 10^-3^) and are slightly more symmetric for average gene expression levels (asymmetric *vs.* symmetric jump branches: 48.58% *vs.* 51.42%, two-sample test for equality of proportions, χ^2^test, *P* = 1.57 × 10^-7^). Our results were robust after changing the branch jump posterior threshold from 0.7 to 0.5 (Supplementary Tables S1 and S2).

When we applied PICs on the phylogenies supporting trait jumps, we observed a clear support for the ortholog conjecture, with more than 21% higher contrasts for paralogs than orthologs (Supplementary figs. S11A and S11B). This suggests that higher PICs could sometimes be due to these rapid shifts in trait values. When we removed the contrasts of the nodes for which the daughter branch(es) experience rapid shifts in their traits, the ortholog conjecture is no longer supported for the traits (Supplementary figs. S11C and S11D). This confirms that trait jumps play an important role in duplicate evolution. Interestingly, there is also support for the ortholog conjecture for the phylogenies not supporting a trait jump model (Supplementary figs. S11E and S11F), implying that these are genes which evolve after duplication in a more continuous manner. Our results were robust when we replicated our analyses on teleosts only (Fig. 5; Supplementary fig. S12). Of note, the nodes with high support for jumps are not always the same as nodes with high PICs (Supplementary fig. S13). Elevated contrasts and high probability of trait jumps thus do not describe the same process.

**Figure 5:**
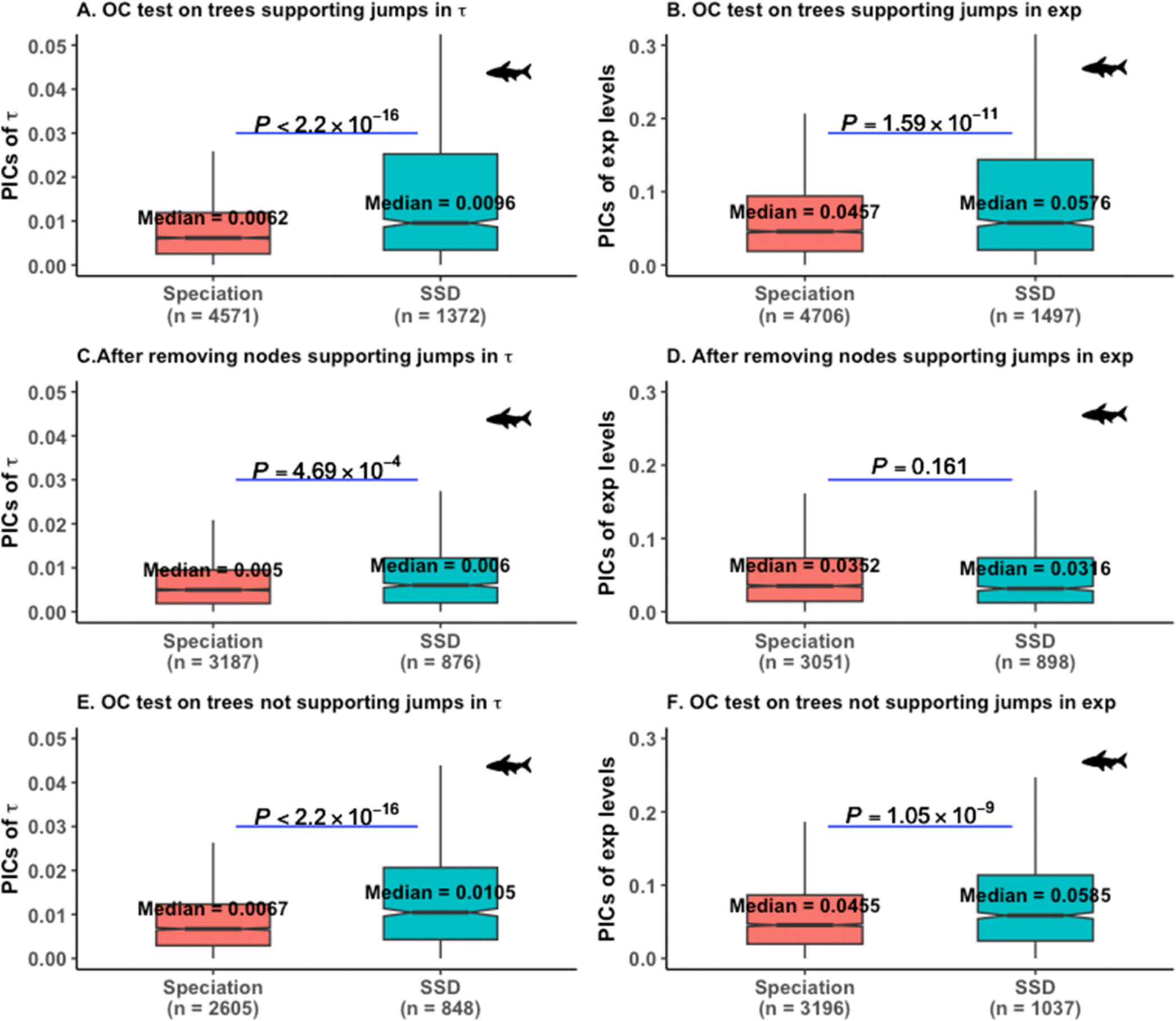
Trait jump test of the ortholog conjecture on small-scale duplicate trees in teleosts. We used a strict branch jump posterior cutoff of ≥ 0.7 as an evidence of trait jump. Only teleosts-specific clades were used for the analyses. The ortholog conjecture tests on (A) 1557 calibrated contrasts standardized strict SSD trees supporting rapid jumps in τ, and on (B) 1593 strict SSD trees supporting jumps in average gene expression levels with the above-mentioned threshold. (C)-The ortholog conjecture tests after removing contrasts of nodes whose daughter branch(es) experienced jumps in the corresponding trait for 1557 and 1593 strict SSD trees, respectively. (E)-(F) The ortholog conjecture tests on 925 and 1124 strict SSD trees for which the trait jump model is not supported for τ and for average gene expression levels by any of their branches, respectively. *P* values are from Wilcoxon two-tailed tests.

### Tissue-specific trait jumps

We next performed our analyses separately on each tissue to investigate whether jumps in expression following duplication were tissue-specific or shared between tissues. Since the trait jump model was supported for SSD and speciation branches, but not genome duplication branches, in the global analyses (Figs. 4 and 5; Supplementary figs. S11, S12 and S14), we restricted the tissue-level analysis to strict SSD trees. Most of these trees supported the trait jump model for all 6 tissues (Fig. 6).

**Figure 6:**
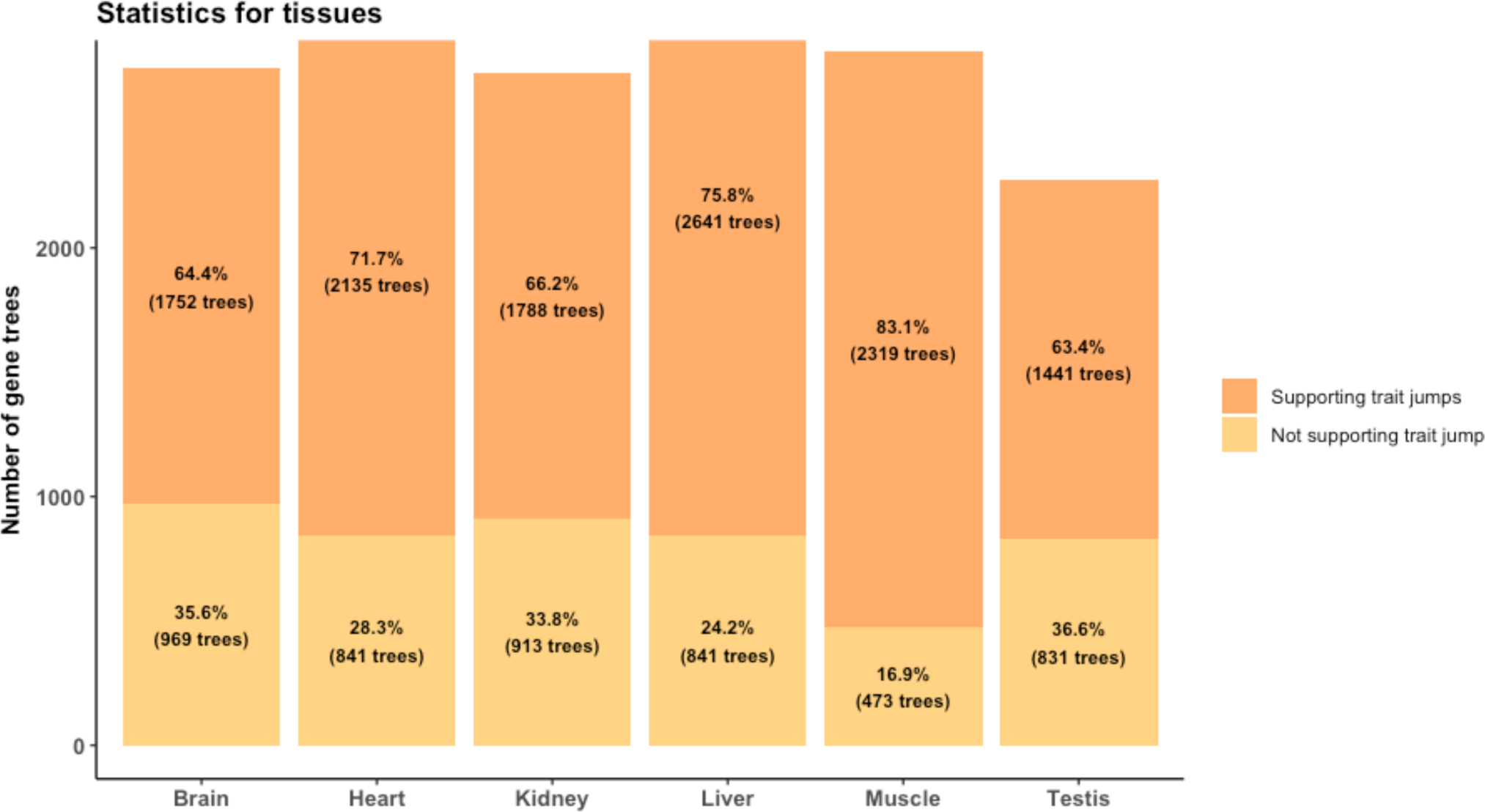
Statistics of contrast standardized small-scale duplicate trees supporting jumps in each tissue. Only strict SSD trees were used. For each tissue, there are more trees supporting trait jumps than not (two-sample test for equality of proportions, χ^2^test, P < 2.2 × 10^-16^).

When we applied our PICs on the trusted SSD phylogenies supporting jumps in their traits for individual tissues, we observed a strong support for the ortholog conjecture in brain, kidney and testis (Supplementary figs. S15-S20). There is also support for the ortholog conjecture in heart, but only in the teleost-only analysis (Supplementary fig. S16). Removal of nodes for which one or of both the daughter branches experiences rapid shifts in their gene expression traits removes the signal of the ortholog conjecture for these four tissues, including teleost-only heart (Supplementary figs. S15C-S15F, S16C, S16F, S17C-S17F, S20C-S20F).

### Relation between coding sequence positive selection and evolutionary shifts in expression

To investigate a potential role of positive selection on genes which shift expression after duplication, we turned to the detection of positively selected sites in the protein coding sequence. We aim to test whether positive selection on the protein is associated with rapid shifts in expression of small-scale duplicates.

Parsing the positive selection data for vertebrates from the Selectome database (Moretti et al. 2014) and applying a false-discovery-rate correction (Storey and Tibshirani 2003; Anisimova and Yang 2007; Studer et al. 2009) cutoff of q < 0.10, we found less than 100 cases with both protein-level positive selection and expression jumps. To gain power, we used the sum of log-likelihood ratios (Daub et al. 2017), where higher log-likelihood ratio (ΔlnL) score indicates that the data have more support for positive selection relative to a nearly-neutral null model. Summing these scores provides the overall support for positive selection over a set of genes. Branches supporting rapid shifts in expression traits have indeed higher ΔlnL scores for paralogs than for orthologs, thus more evidence for positive selection (Supplementary figs. S21A and S21B). While the highest ΔlnL scores were for paralogs with expression shifts, removal of branches supporting rapid jumps in gene expression traits still showed higher ΔlnL scores for paralogs (Supplementary figs. S21C and S21D). Our trends remain mostly unchanged when we repeated our analyses for teleosts only (Fig. 7), with smaller sample sizes. Thus there is overall more positive selection on protein-coding sequences after duplication than speciation, and even more for genes which shift in expression. This supports the ortholog conjecture regarding protein sequence evolutionary rates, as observed by David et al. (2020) and Fukushima and Pollock (2020).

**Figure 7:**
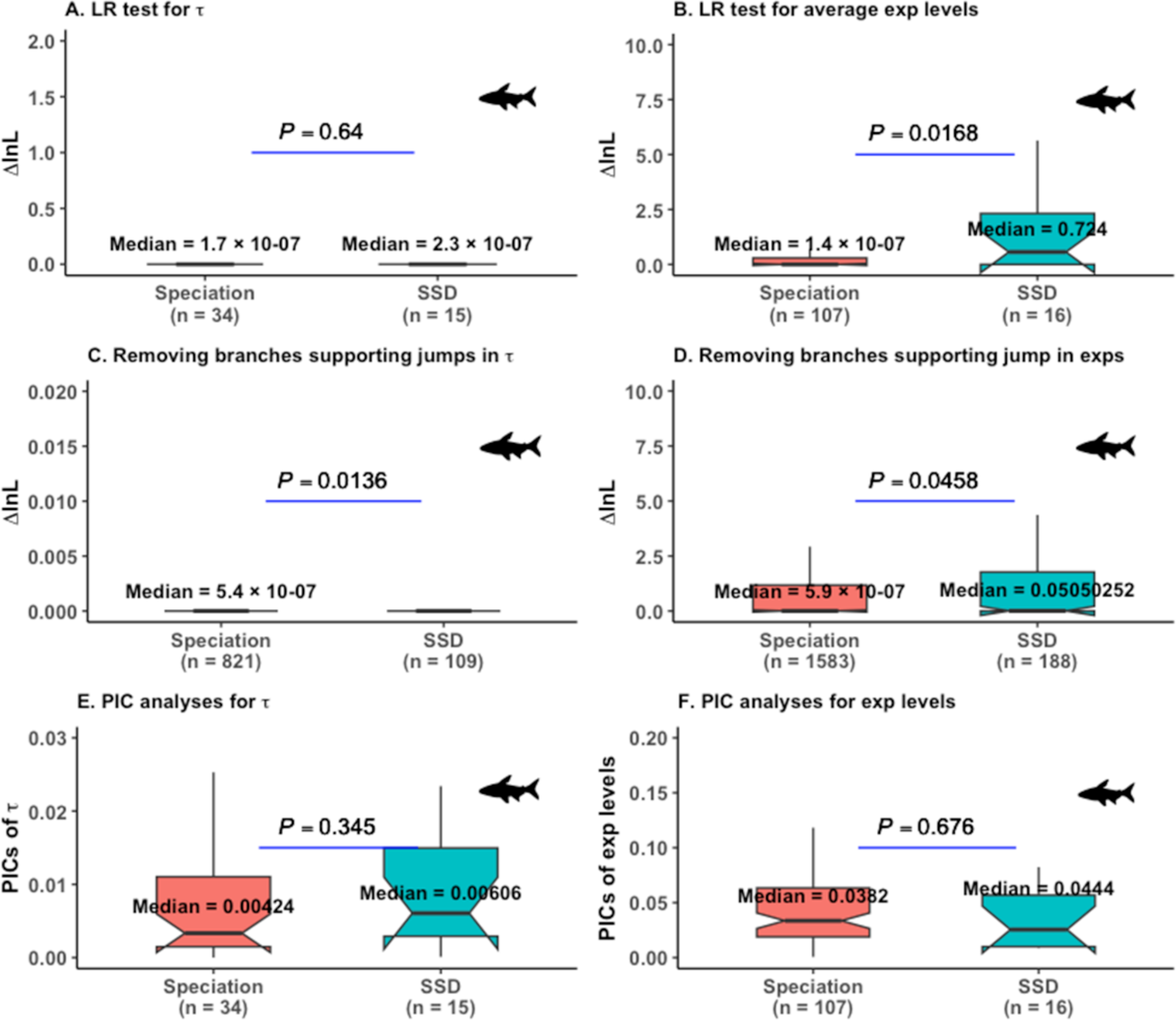
Positive selection and trait divergence analyses for contrast standardized strict SSD trees in 5 teleost fishes. We only used phylogenies supporting an evolutionary trait jump, and then considered teleosts-specific branches for these analyses. We used a strict posterior probability cutoff of ≥ 0.7 to identify branches with rapid jumps in traits. exps: Gene expression levels, PICs: phylogenetic independent contrasts, LRT: likelihood ratio test.

## Discussion

Understanding the dynamics of gene function change after duplication is important both to our fundamental knowledge of gene evolution, and to the practical use of the ortholog conjecture (Koonin 2005; Studer and Robinson-Rechavi 2009; Nehrt et al. 2011; Altenhoff et al. 2012; Chen and Zhang 2012; Gabaldón and Koonin 2013; Rogozin et al. 2014; Kryuchkova-Mostacci and Robinson-Rechavi 2016; Dunn et al. 2018; Fukushima and Pollock 2020; Stamboulian et al. 2020; Begum and Robinson-Rechavi 2021). Indeed, the function of the majority of proteins is not experimentally determined, but rather is extrapolated from homologs, often specifically from orthologs. Although the ortholog conjecture remains debated (Nehrt et al. 2011; Dunn et al. 2018; Stamboulian et al. 2020), we previously found some support for the conjecture for a single trait, tissue specificity τ, from eight vertebrate species using a properly controlled phylogenetic independent contrasts method, as well as using phylogenetic modeling (Begum and Robinson-Rechavi 2021). In that study we did not distinguish types of duplication, and only tested for branch-specific changes in gradual rates.

Here we provide strong confirmation for the ortholog conjecture on gene expression in vertebrates: it holds both for average expression and for tissue-specificity; it holds with a use of phylogenetic independent contrasts which accounts for the long-term impact of duplication on the next speciation branch; and it holds for both small-scale and whole genome duplicates. Our results are in agreement with the results of pairwise-based analyses (Altenhoff et al. 2012; Chen and Zhang 2012; Gabaldón and Koonin 2013; Rogozin et al. 2014; Kryuchkova-Mostacci and Robinson-Rechavi 2016). While the usual interpretation of the orthologue conjecture has considered a change in sequence or trait evolutionary rates over a phylogenetic branch, there could also be sudden changes of trait values without altering the evolutionary rates (Begum et al. 2021). And indeed, we find evidence for such a rapid trait jump model for evolution after duplication.

Evolutionary jumps have been shown to affect the evolution of traits in adaptive landscapes (Simpson 1944; Bokma 2008; Landis et al. 2013; Duchen et al. 2017; Duchen et al. 2021; Gao and Wu 2022), but have been rarely applied to the evolution of gene function. Yet small-scale gene duplication can be a rapid and potentially asymmetric phenomenon. Indeed, we find evidence for the existence of a trait jump mechanism for small-scale duplicates. This can correspond to a sudden change of function, e.g., from incomplete duplication of the coding portion of the progenitor locus or of its regulatory regions or a change in chromatin environment (Lynch and Katju 2004; Dennis et al. 2012; Dougherty et al. 2017; Guschanski et al. 2017). Paralogs can differ in structure from their progenitor copy (Dougherty et al. 2017), and can acquire novel promoters (e.g., gene fusion), or partially or completely lose regulatory elements (e.g., RNA-mediated duplication, Kaessmann et al. 2009). These processes can lead to rapid functional changes, notably through altered expression patterns. These mechanisms rapidly change the trait values rather than changing their evolutionary rates over long evolutionary time. Tissue-specific support for trait jumps after duplication appears to differ between organs, with strongest support in brain, testis and kidney (Figs S15-S20). On the other hand, genome duplication does not seem to lead to sudden or very asymmetric changes in gene expression, as shown by the lack of support for the jump model, which is consistent with the short-term conservation of regulatory environment (Figs. 2 and 3, Supplementary figs. S4 and S9). Interestingly, even for small-scale duplicates, those which do not fit a jump model still support the ortholog conjecture, i.e. with a change in gradual evolutionary rate. Thus the two models of divergence after duplication co-exist: rapid jumps after some small-scale duplications, and gradual changes after whole genome and other small-scale duplications.

Positive selection on the coding sequence can contribute to the evolution of gene function following duplication (Han et al. 2009; Pegueroles et al. 2013). We found indeed in general higher evidence for positive selection after duplication. We also find slightly more evidence for positive selection on the branches supporting rapid jumps in their traits following duplication than speciation, but tests are not significant. This might be in part due to a lack of power, as sample sizes for genes with evidence of both processes are small. Thus we cannot exclude a relation, but if there is one it is weak and would need larger sample sizes to detect reliably.

In conclusion, we find strong support for rapid evolution of gene expressions after duplication, both by sudden jumps and by long-term acceleration of evolutionary rates. Processes differ between small-scale and genome-scale duplications, but in all cases support the ortholog conjecture, i.e. stronger divergence of paralogs than of orthologs. Thus duplication is critical to the evolution of genome function, and is best studied with a careful use of phylogenetic methods.

## Materials and Methods

### Processing gene trees

43491 empirical gene trees were parsed from Genomicus v.95.01 (Nguyen et al. 2022). This version includes the division of some large trees (> ∼400 homologs) into subtrees, like Ensembl in later versions (Martin et al. 2023). Our analyses should not be affected by this, since the missing nodes in the split trees are mainly first round or second round WGD nodes at the base of vertebrates (Parey et al. 2020), which are not the focus of this study. Trees were processed to annotate “speciation”, and “duplication” to the internal node events using available information from the tree data object (“D” = 0 as “speciation” and “D” = “Y” as “duplication”). We estimated tissue specificity, τ, and average gene expression levels from ten tetrapods (human, mouse, rat, macaque, cow, dog, ferret, opossum, rabbit, and chicken) and from five teleost fishes (zebrafish, medaka, cavefish, northern pike, and tilapia) across six common tissues (brain, heart, kidney, liver, muscle, and testis), and added those estimates to the tree data objects according to their corresponding Ensembl gene ID. After pruning the gene trees to remove tips with missing trait estimates we obtained 17103 trees. We excluded 148 gene trees with only duplication nodes and 7916 gene trees with no duplication nodes, and retained 9039 trees with at least one speciation and one duplication node for further processing. Cross-species normalized gene expression levels (see below) were then added to the tree data object, and all 9039 trees were retained after filtering for the eventuality of trees with genes that were not expressed in any of the six tissues or which had less than four tips.

The gene trees were then time calibrated using speciation clade ages from Timetree (Hedges et al. 2006). For time calibration, we used the “correlated” model of chronos() function from the R “ape” package (Paradis et al. 2004; Dunn et al. 2018; Begum and Robinson-Rechavi 2021). Since the focus of the study was teleost species, we considered tetrapods as the outgroups. We, therefore, excluded trees with no teleost tip data for our analyses. We thus selected 6923 trees for which at least three tips belonged to teleosts, and at least one tip contained trait data for an outgroup tetrapod species.

### Estimation of phylogenetic independent contrasts (PICs)

For each individual trait, phylogenetic independent contrast or PIC was estimated for each internal node of a gene tree using the pic() function of the R “ape” package (Paradis et al. 2004). PIC of a parental node is an absolute ratio of changes in trait values for descendant nodes to the lengths of its two descendant branches, *i.e.*, the expected variance (Felsenstein 1985; Dunn et al. 2018; Begum and Robinson-Rechavi 2021), in millions of years (My).

### Classification of gene trees based on types of duplications

We identified synteny validated teleost-specific third round (relative to previous two rounds at the base of vertebrates) whole genome duplicates (FishWGDs), or ohnologs, from the database OHNOLOGS v.2 (Singh and Isambert 2020). Out of five teleosts, we obtained 3R ohnolog pairs only for two (*e.g.*, zebrafish and medaka) from this database. We applied relaxed criteria (q-score (outgroups) < 0.05 and q-score (self-comparison) < 0.3) as available in OHNOLOGS database (Singh and Isambert 2020) to obtain a large dataset of ohnologs. Only “protein_coding” genes with duplication time annotated as “FishWGD” were used. We annotated the most recent common ancestral node of the identified ohnolog pairs in each gene tree as “FishWGD”. Other duplication nodes in the same trees were annotated as small-scale gene duplication or “SSD”.

The teleost fish underwent a third round of whole genome duplication at the base of the Osteoglossocephalai or the Clupeocephala clade, which has been dated 225 − 333 My ago (Berthelot et al. 2014; Parey et al. 2020; Parey et al. 2022). This ambiguity in the position of the teleost WGD is presumably because Genomicus v.95 trees were based on a different phylogeny than that recently resolved in Parey et al. (2023). We thus considered “FishWGD” annotated to Osteoglossocephalai or Clupeocephala clade descendants as “confident” FishWGD ohnologs. We called the 1159 phylogenies with such confident ohnologs “strict WGD” trees. Other ohnolog-annotated genes of the OHNOLOGS database were classified as “dubious 3RWGDs” when their duplication time did not match with “FishWGD” or when the “FishWGD” annotation matched with taxonomic levels other than Osteoglossocephalai or Clupeocephala. We thus obtained 628 “dubious WGD” trees.

To identify small scale duplication (SSD) trees, we first excluded the 1787 strict or dubious WGD gene trees. From the remaining 5136 gene trees, we then eliminated phylogenies where duplication was present at the Clupeocephala or Osteoglossocephalai taxonomic level. We called those 4139 trees “strict SSD” trees. The other 997 phylogenies were called “dubious SSD” trees, where duplication occurred at the same Clupeocephala or Osteoglossocephalai clade as the 3R WGD. To perform our phylogenetic analysis on dubious SSD trees, we removed contrasts (PICs) of those Clupeocephala and Osteoglossocephalai clades.

### Selection of contrasts standardized trees

Since the performance of the PIC method primarily depends on the assumption of Brownian model (BM) of trait evolution, we need to identify phylogenies for which contrasts (PICs) are adequately standardized according to BM (Felsenstein 1985; Garland 1992; Diaz-Uriarte and Garland 1996; Díaz-Uriarte and Garland 1998; Freckleton and Harvey 2006; Cooper et al. 2016; Begum and Robinson-Rechavi 2021). Node contrasts (PICs) for those trees should be independent of internal phylogenetic parameters, *i.e.*, logarithm of node age, expected standard deviations and node depth in this case. We used diagnostic tests on phylogenies with duplications to draw unbiased inference by PIC method, as recommended earlier (Garland 1992; Diaz-Uriarte and Garland 1996; Díaz-Uriarte and Garland 1998; Freckleton and Harvey 2006; Cooper et al. 2016; Begum and Robinson-Rechavi 2021). In these tests, we perform correlation between the absolute values of PICs and each of the three internal phylogenetic parameters individually for each trait per phylogeny. Any significant correlation (*P* < 0.05) indicates departure from BM, *i.e.*, phylogenetic dependence for that tree. By eliminating trees showing a significant trend (positive or negative), we considered sets of trees passing all three diagnostic tests for further analyses for each trait of our interest. We estimated node age and node depth using branching.times() and node.depth() functions from the R “ape” package, respectively, for each tree (Paradis et al. 2004). We applied the crunch() function of the caper package to perform diagnostic tests (Purvis and Rambaut 1995; Cooper et al. 2016; Orme 2018; Begum and Robinson-Rechavi 2021).

Following this, 746 strict FishWGD trees and 2482 strict SSD trees passed diagnostic tests for τ, and 774 strict FishWGD trees and 2717 strict SSD trees passed diagnostic tests for average gene expression levels. Overall, we obtained 4247 and 4603 trees passing diagnostic tests for τ and average gene expression levels, respectively, including “dubious” FishWGD or SSD trees.

### Randomization test

To confirm biological effects vs. biases in data structure, we used randomization test of trait values on different sets of contrasts standardized trees, following Begum and Robinson-Rechavi (2021).

To randomize, we permuted actual trait values of the tips without changing internal node events for each tree. After randomization, we recalculated the node contrasts (PICs) for each tree using permuted trait values of tips following above stated procedure.

### Expression data

We retrieved processed adult stage paired-end RNA-seq data for seven tetrapods (human: GSE30611, mouse: GSE41367, rat: GSE41367, macaque: GSE41367, cow: GSE41367, opossum: GSE30352, and chicken: GSE41367) and one teleost fish (zebrafish: SRP044781) from Bgee database v.14 (Bastian et al. 2021). Pre-processed expression data for three other tetrapods (dog, ferret, and rabbit) submitted to the NCBI Gene Expression Omnibus (GEO; http://www.ncbi.nlm.nih.gov/geo/) by Chen et al. (Chen et al. 2019) under accession number GSE106077 were directly downloaded from https://portals.broadinstitute.org/evee/. Paired-end Illumina RNA-seq reads of three teleosts (medaka, cavefish, and northern pike) were retrieved from the PhyloFish database (Pasquier et al. 2016). All the tissue data of PhyloFish came from mature stage, except for testis, which came either from adult, or from unknown stage. We fetched paired-end RNA-seq raw reads for tilapia fish directly from NCBI using accession number SRR391690. Developmental stage information was unavailable for dog, ferret, rabbit, and tilapia.

Quality assessment of raw and trimmed reads was conducted using FastQC v.0.11.5 (Andrew 2019). Quality filtration was performed using Trimmomatic v.0.36 (Bolger et al. 2014) to discard adaptor sequence or low-quality reads (settings: LEADING:4 TRAILING:4 SLIDINGWINDOW:4:20 HEADCROP:12 MINLEN:25). Transcript information was downloaded from Ensemble v.95 (Martin et al. 2023) and was used as reference for fast and accurate mapping against the cleaned reads with the aid of kallisto v.0.43.1 (Bray et al. 2016) pseudoaligner. For rapid RNA-seq quantification, we applied quant argument of kallisto. We used transcript abundances in transcripts per million as an estimate of gene expression levels, with log transformation (log_2_TPM). In case of tilapia, we had three biological replicates for each tissue sample. We took average TPM for all the replicates against each tissue to obtain per tissue gene expression level for each gene. TPM values from several sub parts of the tissues for tetrapods from Bgee database (Bastian et al. 2021) are averaged to obtain gene expression levels of the main organ or tissue. Teleosts specifically have an organ called “head kidney”, which comprises about 20% of the anterior part of the kidney. Considering it as “kidney”, we obtained six common tissues (brain, heart, kidney, liver, muscle, and testis) for all the vertebrates of this study.

### Normalization of gene expression data

For a few species of this study, the developmental stages of tissues were unknown (see above). Most PhyloFish samples, except testis, were extracted from females (Pasquier et al. 2016). On the other hand, data from Bgee (Bastian et al. 2021) sometimes came from male, sometimes from female, or also from mixed sex in some cases (e.g., heart and kidney expression of zebrafish). In addition, these data come from different studies and different species. Thus we needed to normalize gene expression levels to compare them across species and tissues.

For cross-species expression data normalization, we adopted a scaling procedure following Brawand et al. (2011). Among the 1077 1:1 orthologs of all vertebrate genes with their expression values in the interquartile range, we identified the top 500 conserved genes that have the maximum conserved ranks among tissues. Median expression levels for those 500 genes were assessed in each tissue for each species. We then derived scaling factors for each tissue of a species using median of human tissues as references. Finally, we applied those scaling factors on the log transformed corresponding tissue dataset of 15 species to obtain normalized gene expression level for each gene of a species in that tissue.

### Estimation of trait data

Tissue specificity, τ, of a gene was computed across all the six tissues of each species using the following equation (Yanai et al. 2005):

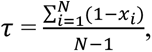

where N is the total number of tissues or organs and *x_i_* is the log_2_(TPM) value of gene x in i^th^ tissue. τ close to 1 indicates higher tissue specific function of a gene, while more ubiquitous expression is denoted by values close to 0 (Yanai et al. 2005; Kryuchkova-Mostacci and Robinson-Rechavi 2016).

Average gene expression level was estimated by taking the mean of cross species scaling factor normalized log_2_(TPM) values across all the six tissues for each gene per species.

### Estimation of rapid trait jump

Evolutionary jumps were estimated with the software “levolution” (Duchen et al. 2017). The input consists of an ultrametric tree in Newick format, and a two-column file with trait data for each tip of the tree. In our case the trees used were the gene trees described above, and the trait data were the normalized gene expression for the different tissues, as well as for the τ and average gene expression levels. “levolution” was then run separately for individual trait on contrast standardized gene trees.

Briefly, “levolution” works by calculating the variance-covariance matrix of the traits, and assigning a starting vector of jump configurations at every branch of the tree. Through a Markov chain Monte Carlo (MCMC) method, jump configurations are then updated one branch at the time with fixed hierarchical parameters, such as the jump rate, Brownian rate, jump strength, and root state. Two steps take place during the MCMC run. First, an Expectation-Maximization approach is used to infer maximum likelihood (ML) estimates of these hierarchical parameters. Then, the position of jumps along the tree is inferred by using an empirical Bayes approach, where the hierarchical parameters are fixed to their ML estimate, and the MCMC is then used to calculate posterior probabilities of jumps at a particular branch (Duchen et al. 2017).

Mathematically, this jump model is depicted as follows. Given ***x*** as the set of traits at the tips of the tree, the likelihood of ***x*** given the hierarchical parameters is f(x|hierarchical parameters) = 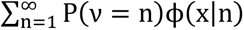, where ***n*** is the number of jumps, *P*(*v* = *n*) is the Poisson probability of observing ***n*** jumps, and *φ*(*x*|*n*) is the multivariate normal probability of observing traits ***x*** given that ***n*** jumps are present (Duchen et al. 2017, Eq. 4). The sum over all possible jump configurations is taken over by the MCMC approach described above, and the entries in the variance covariance matrix in *φ*(*x*|*n*) are given by the sum of the variance present along each branch and the variance due to the jumps.

### Analyses of positive selection

We retrieved precomputed statistics on positive selection from Selectome (Moretti et al. 2014), a database that applies the branch-site likelihood test for selection to Ensembl gene trees (release 98, Martin et al. 2023). Each processed Genomicus tree (See ‘Processing gene trees’) was mapped, if possible, to the corresponding Selectome tree by Ensembl gene IDs. Of 6923 processed Genomicus trees, 6649 trees were successfully mapped to their corresponding tree in Selectome. Because the Ensembl-based Selectome trees sample more taxa than we have included in our study, we retrieved the likelihood-ratio test statistic only for nodes that have matching taxonomic labels between each mapped Selectome and Genomicus tree, and excluded taxa that are not contained in the former.

There were 116 (respectively 402) strict SSD trees supporting both positive selection as well as jumps in τ (respectively in average expression levels), in their branches for the 15 vertebrates. For teleost-specific analyses, there were only 31 and 78 such trees.

### Script and data availability

All the scripts and gene expression data files are available on GitHub: https://github.com/tbegum/Phylogenetic-modeling-of-evolutionary-trait-jumps. For reproducibility, “norm_exp_15spe_4tips_new.RData” file is archived at Zenodo (https://doi.org/10.5281/zenodo.10418895).

### Details of other R packages used

We performed other analyses and plotting in R version 4.0.2 (R Core Team 2018) using rphylopic (Chamberlain 2018), phytools (Revell 2012), treeio (Guangchuang 2018), ggtree (Guangchuang et al. 2017), digest (Antoine Lucas et al. 2018), dplyr (Wickham et al. 2017), ggplot2 (Wickham 2016); gtools (Warnes et al. 2018), gridExtra (Auguie 2017), tidyverse (Wickham 2017), stringr (Wickham 2019), and png (Urbanek 2013) libraries.

## Acknowledgements

We sincerely thank all the members of the Robinson-Rechavi and Salamin groups for their help and useful discussions. Parts of the computations were performed at the Vital-IT Center for high-performance computing of the SIB Swiss Institute of Bioinformatics, as well as using the Wally cluster of the University of Lausanne. Work supported by Swiss National Science Foundation grant 31003A_173048.

## Supporting Information

**Figure S1:**
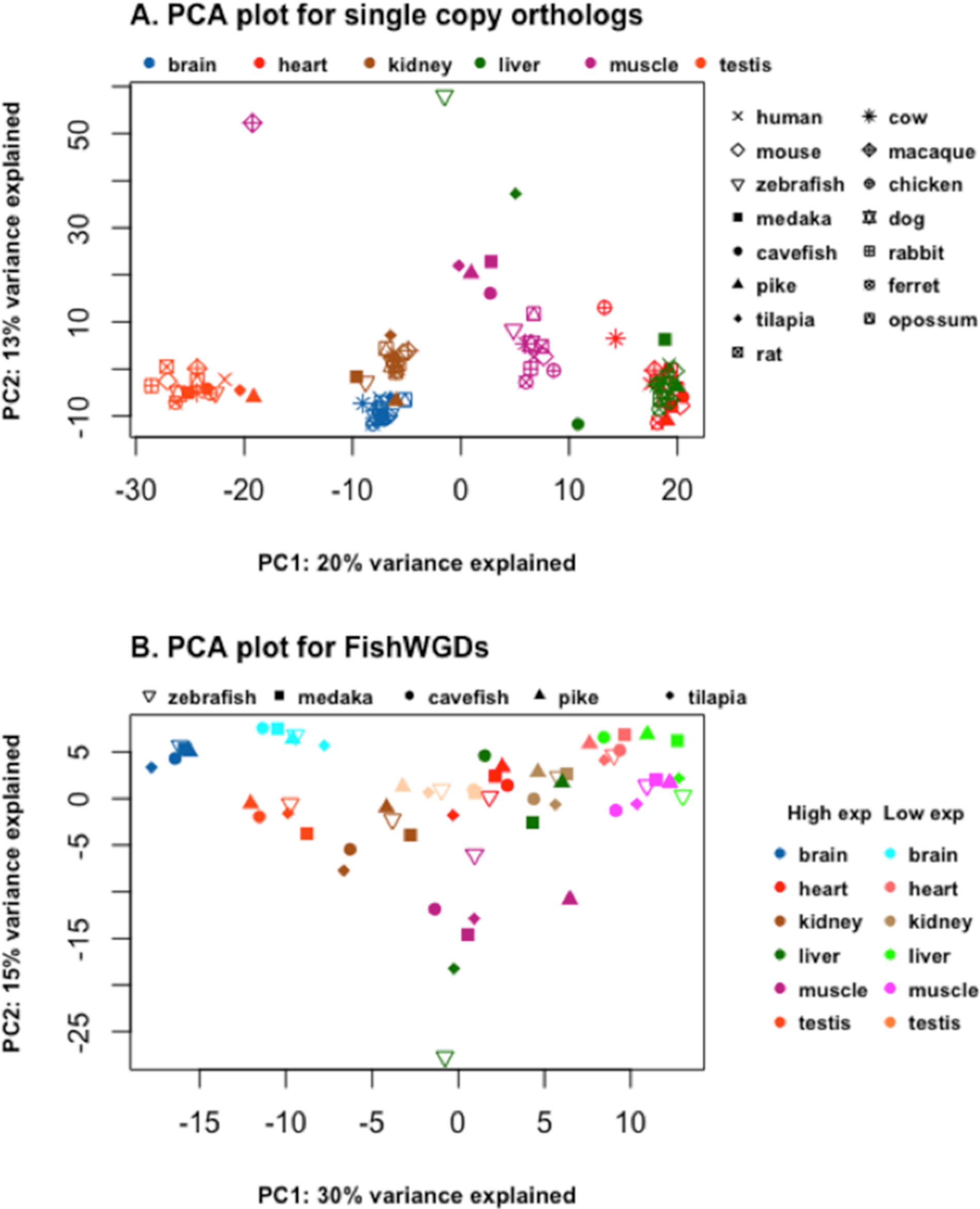
Principal component analysis (PCA) plots. (A) This plot uses normalized expression levels of 1077 1:1 orthologous gene set from 15 species. (B) We consider strict ohnologs or 3R whole genome duplicates (FishWGDs) from 5 teleosts in this plot. Normalized gene expression levels of 300 ohnolog set are used here. Duplicates in each teleost fish are classified based on their expression levels as “highly expressed” and “lowly expressed” ohnologs. Tissue colors for the highly expressed ohnolog copy and the lowly expressed ohnolog copy for a species are labeled as “High exp” and “Low exp”. The plot shows that highly expressed ohnologs cluster separately from their lowly expressed copy for each tissue. Percentage of variations are indicated in each axis. PC1: Principal component 1, PC2: Principal component 2.

**Figure S2:**
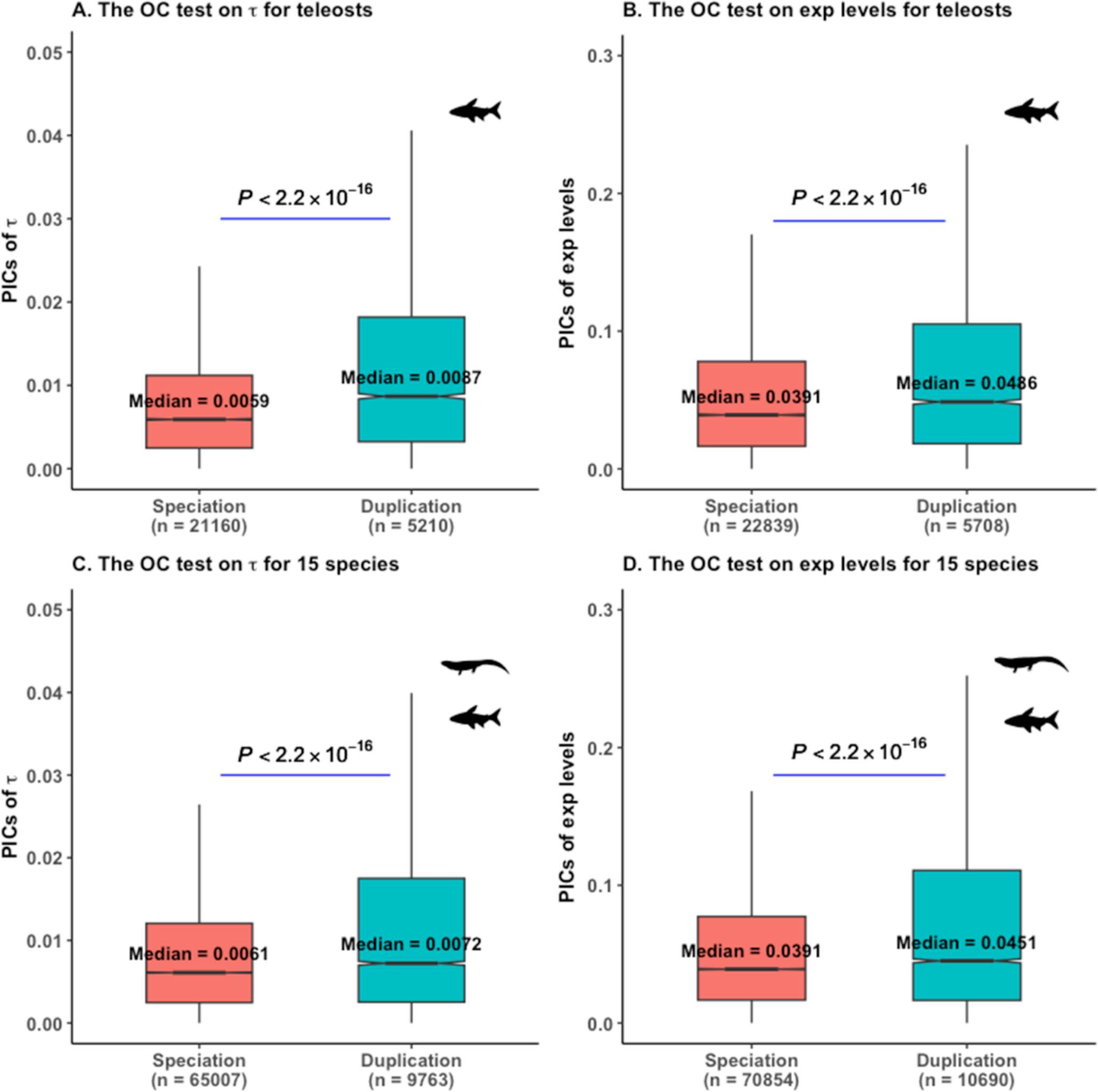
Phylogenetic independent contrast ortholog conjecture (OC) tests. PICs: Phylogenetic independent contrasts, exp: expression. Plots are based on 4247 and 4603 trees for τ and for average gene expression levels, respecively. For (A)-(B), we considered the PICs of teleosts specific speciation and duplication nodes for our analyses, while PICs of all the speciation and duplication nodes are used in (C)-(D). *P* values are from Wilcoxon two-tailed tests.

**Figure S3:**
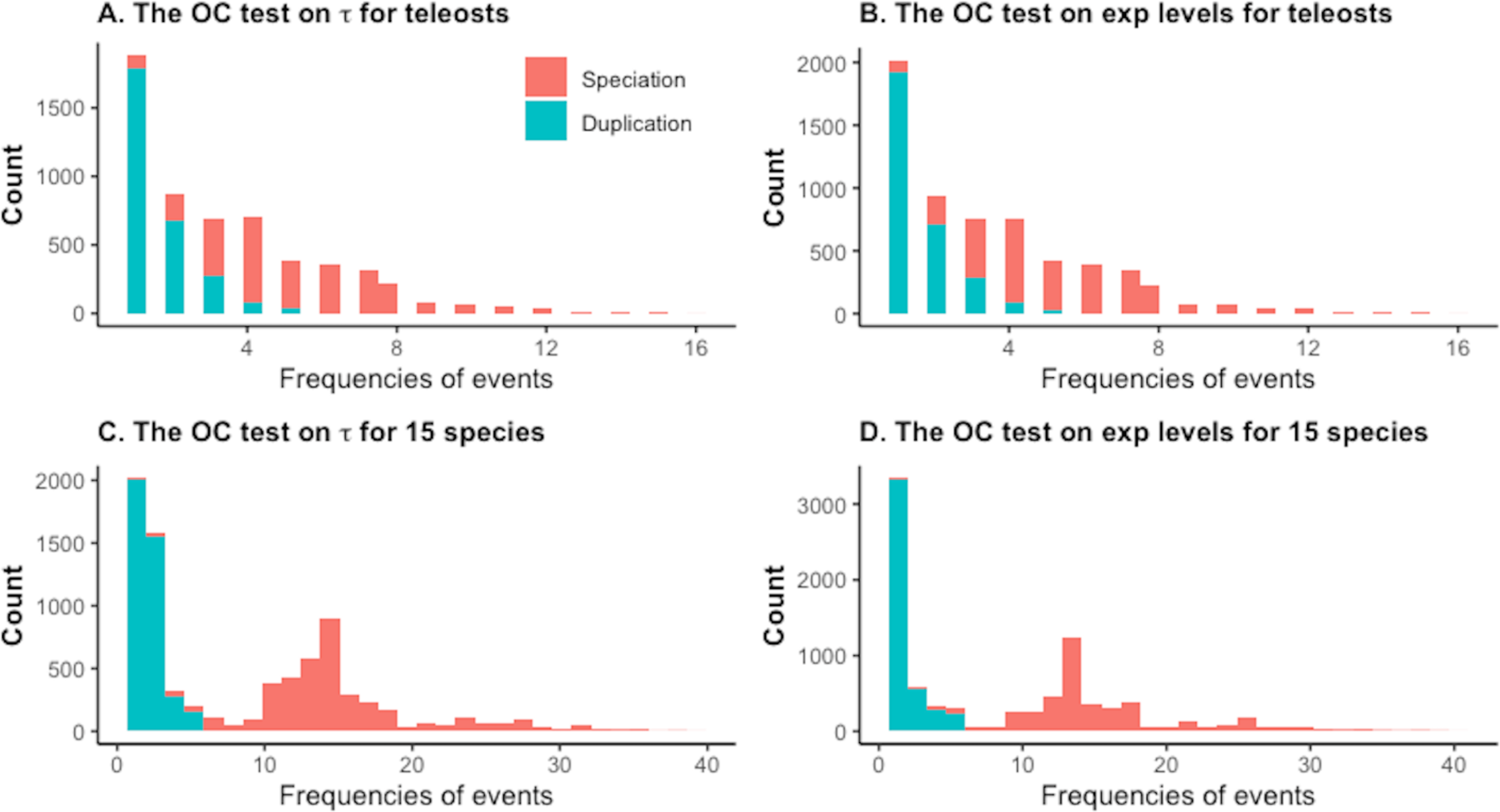
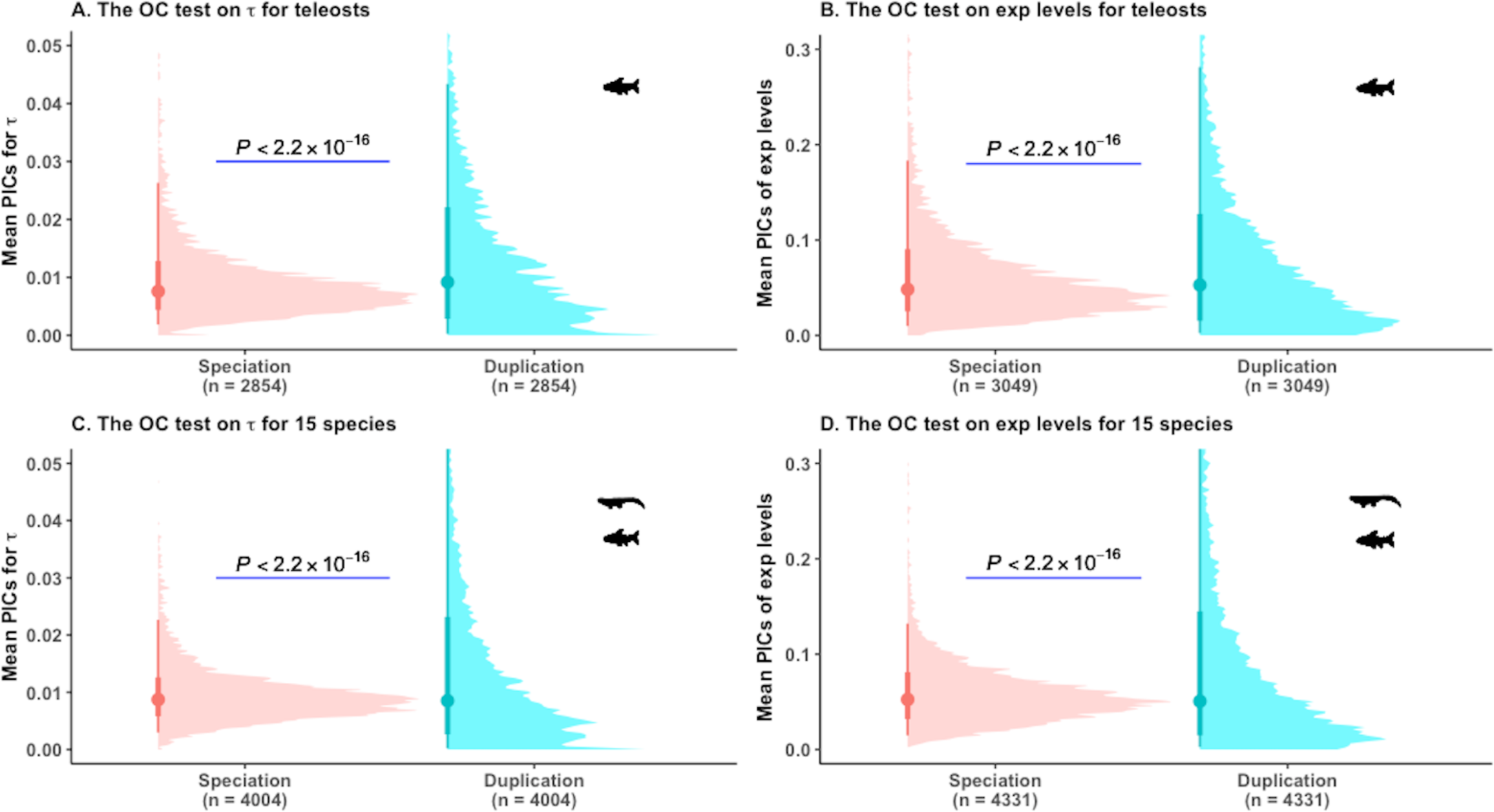
(A) Distributions of events for traits in teleosts and in vertebrates. (B) Phylogenetic independent contrast ortholog conjecture (OC) tests. Instead of using each node of a gene tree as a data point, we used the mean contrast per event for each tree to obtain one datapoint per gene family. Since we obtained one datapoint for speciation and one datapoint for duplication for each tree, paired Wilcoxon test was used to compare the difference. PICs: Phylogenetic independent contrasts, exp: expression. For both (A) and (B), we only used gene trees with ≤5 duplication nodes.

**Figure S4:**
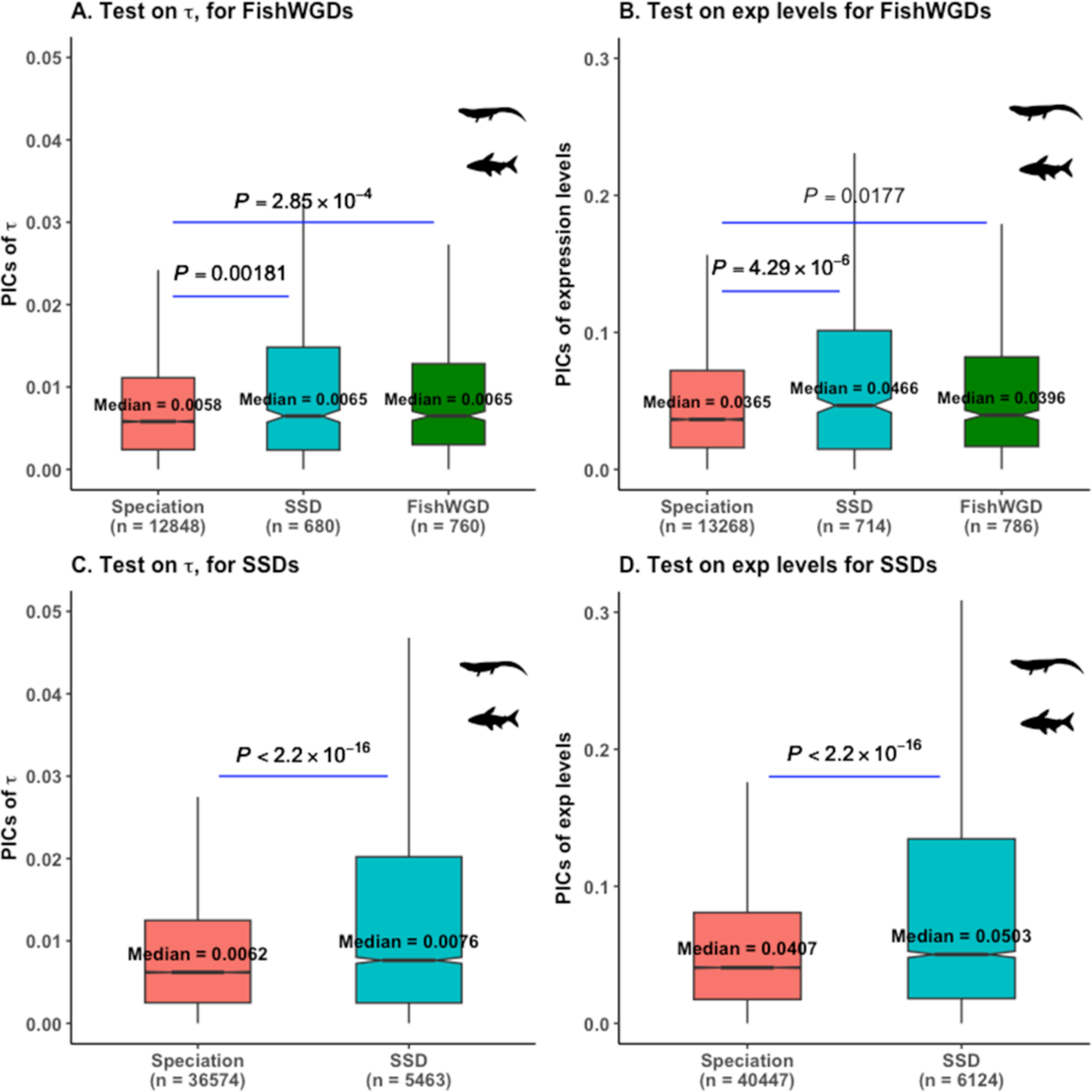
The ortholog conjecture (OC) tests using calibrated contrasts standardized trees in 15 vertebrates. 746 and 774 trees with whole genome duplications were used in (A)-(B), while 2482 and 2717 trees with small-scale gene duplications were used in (C)-(D). PICs: Phylogenetic independent contrasts, FishWGD: Fish specific 3R whole genome duplication, 3RWGDs: ohnologs, SSDs: small-scale duplicates. Duplications in (A)-(B) refers to SSDs like (C)-(D). *P* values are from Wilcoxon two-tailed tests.

**Figure S5:**
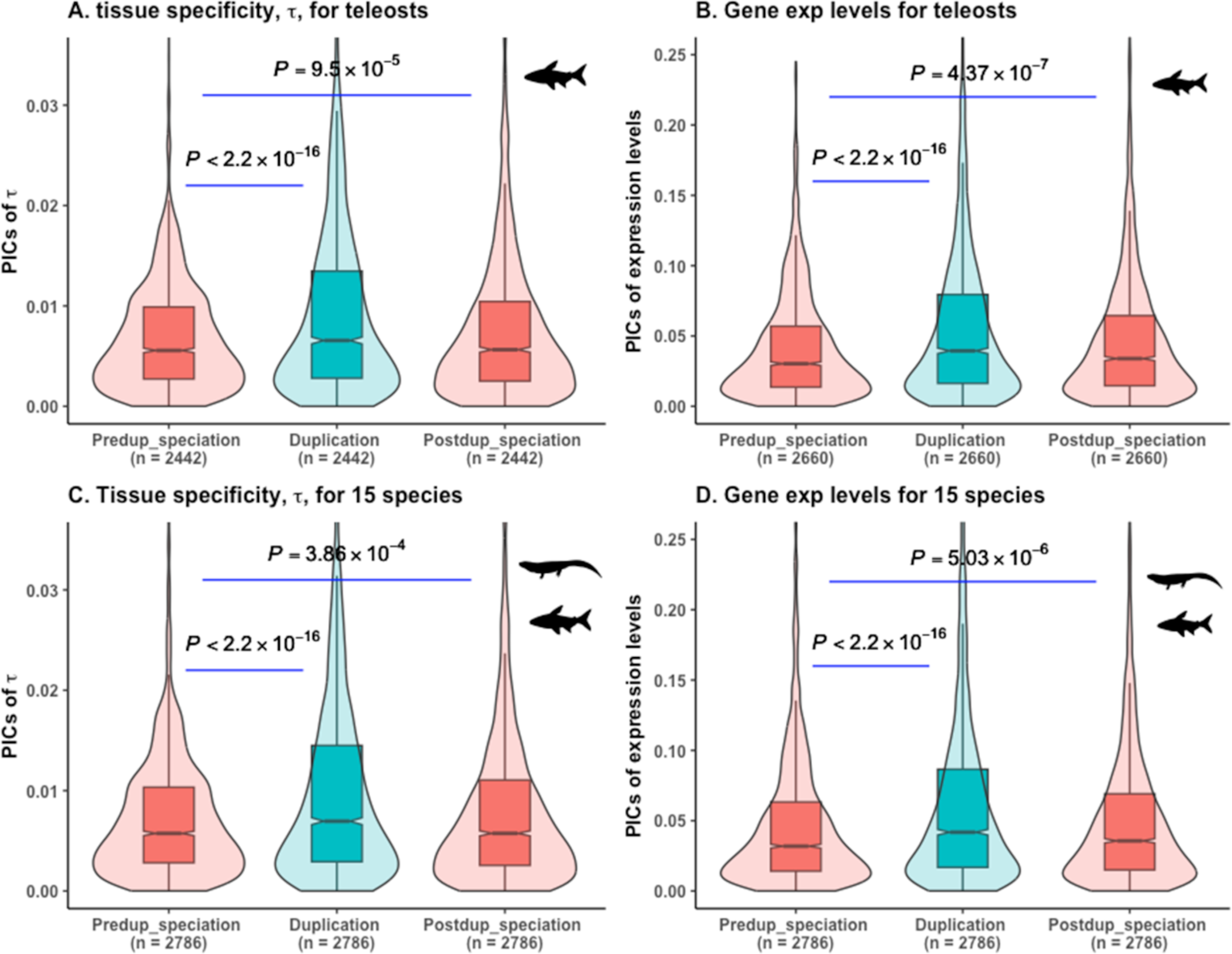
Modified phylogenetic approach to test the ortholog conjecture for teleosts and for vertebrates, respectively. We used the oldest duplication node along a path of a phylogeny for identification of pre-duplication and post-duplication speciations. For (A)-(D) plots are based on 1375 and 1493 trees for τ and for average gene expression levels, respecively. For (A) and (B), we consider the contrasts of teleosts specific duplication and descendant speciation nodes in absence of teleosts specific pre-duplication speciation nodes contrasts. Results of (C) and (D) are based on all 15 vertebrate species. PICs: Phylogenetic Independent Contrasts, exp: expression, Predup_speciation: Pre-duplication speciation, Postdup_speciation: Post-duplication speciation. Since contrasts are estimated for each tree, paired Wilcoxon test is used to compare the difference.

**Figure S6:**
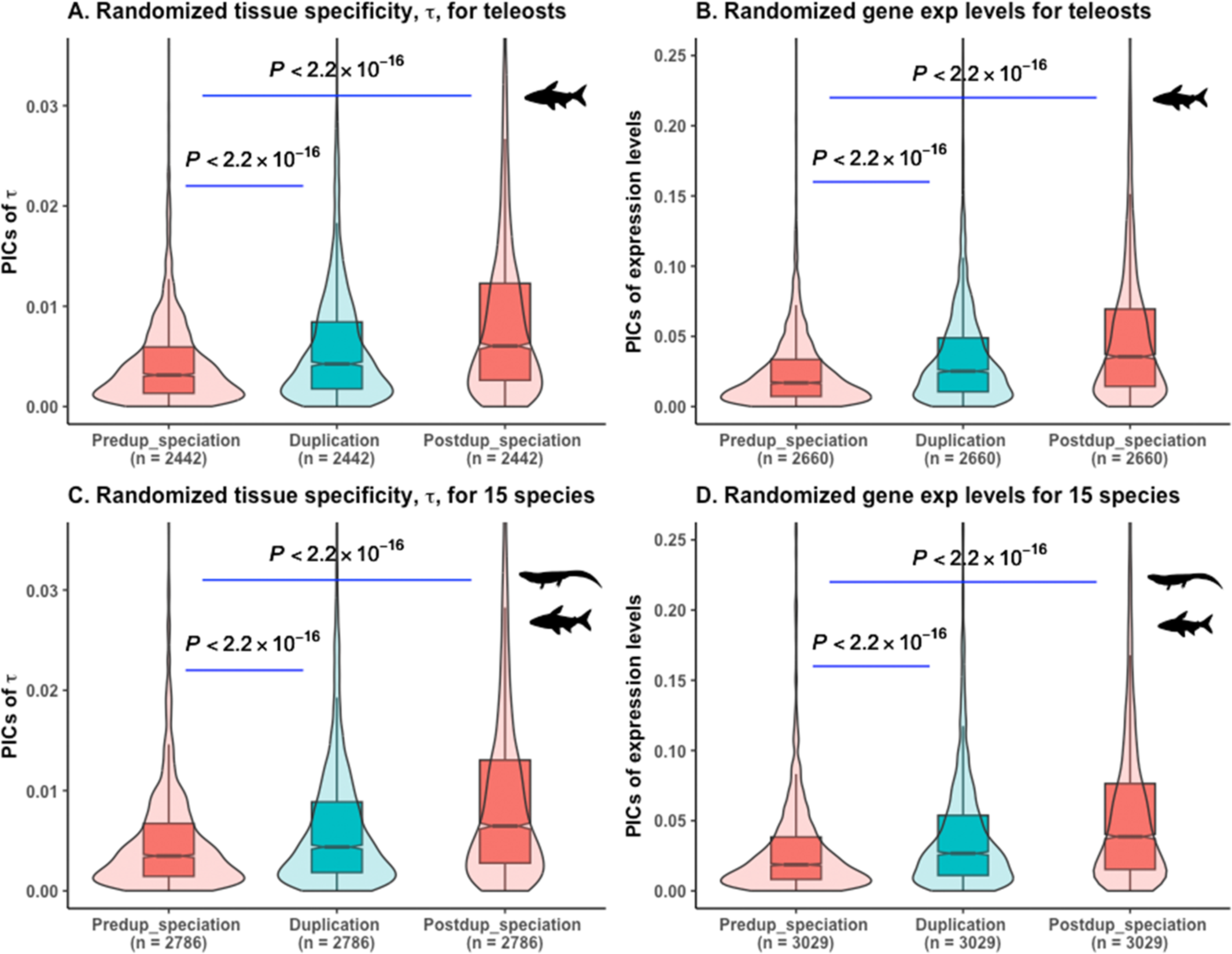
Randomization tests on our modified phylogenetic approach for teleosts and for vertebrates, respectively. Randomization of traits through tip shuffling allow us to use the same nodes used for the ortholog conjecture tests in Fig. S3. For (A)-(D) plots are based on randomization tests on 1375 and 1493 trees for τ and for average gene expression levels, respecively. We considered the contrasts of teleosts specific gene or genome duplication and descendant speciation nodes in (A)-(D) in absence of teleost-specific pre-duplication speciation nodes contrasts. PICs: Phylogenetic Independent Contrasts, exp: expression, Predup_speciation: Pre-duplication speciation, Postdup_speciation: Post-duplication speciation. Since contrasts are estimated for each tree, paired Wilcoxon test is used to compare the difference.

**Figure S7:**
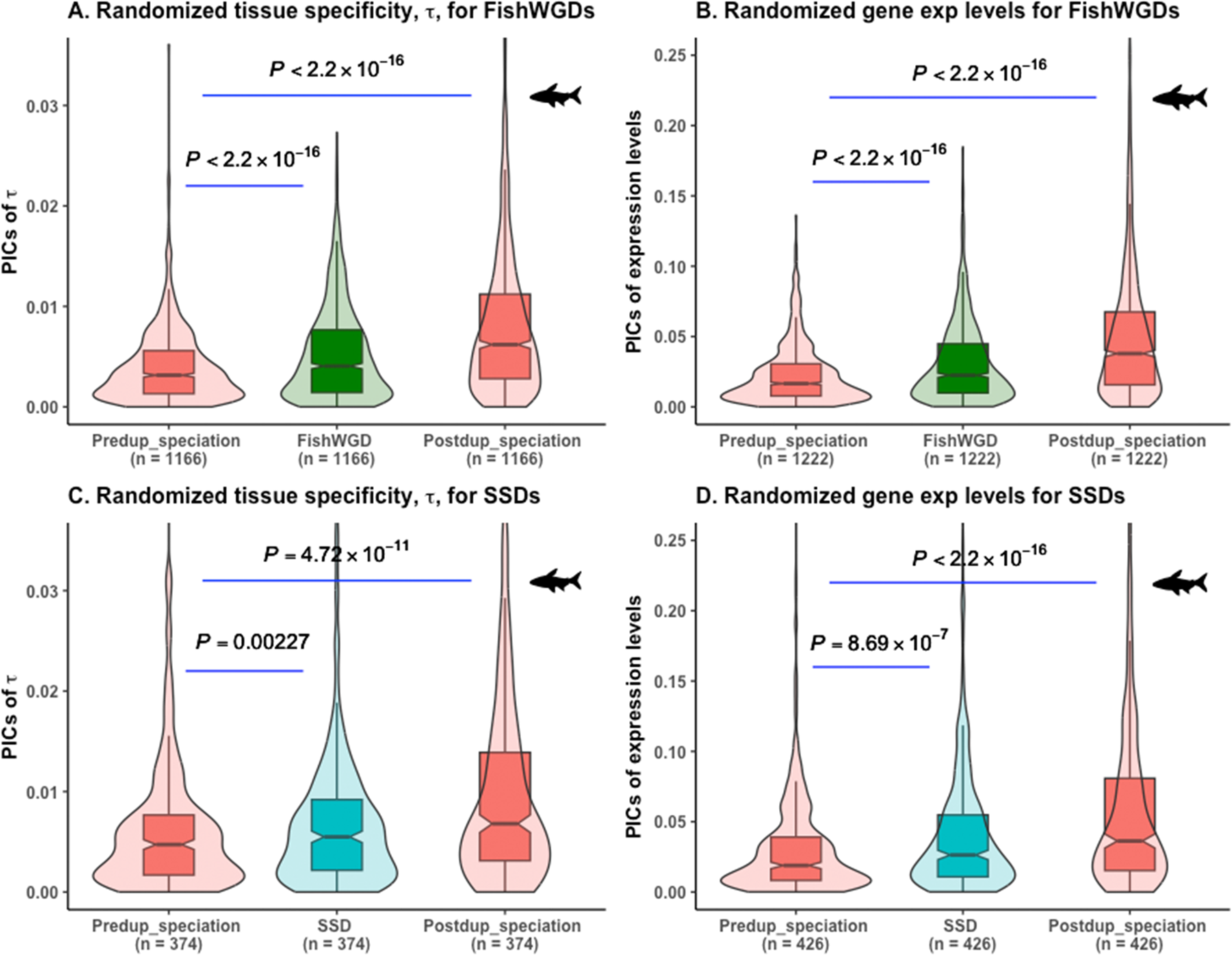
Randomization test on our modified phylogenetic approach for strict 3RWGD and for strict SSD trees in teleosts, respectively. Randomization of traits through tip shuffling allow us to use the same nodes used for the ortholog conjecture tests in Fig. 4. (A)-(D): Plots are based on randomization tests for 581 and 610 strict 3R whole genome duplicate trees and for 339 and 377 strict small-scale duplicate trees for τ and average gene expression levels, respecively. We considered the contrasts of teleosts specific gene or genome duplication and descendant speciation nodes in (A)-(D) in absence of teleost-specific pre-duplication speciation nodes contrasts. PICs: Phylogenetic Independent Contrasts, exp: expression, Predup_speciation: Pre-duplication speciation, Postdup_speciation: Post-duplication speciation, FishWGD: fish specific 3R whole genome duplication, 3RWGDs: ohnologs, SSDs: small-scale duplicates. Since contrasts are estimated for each tree, paired Wilcoxon test is used to compare the difference.

**Figure S8:**
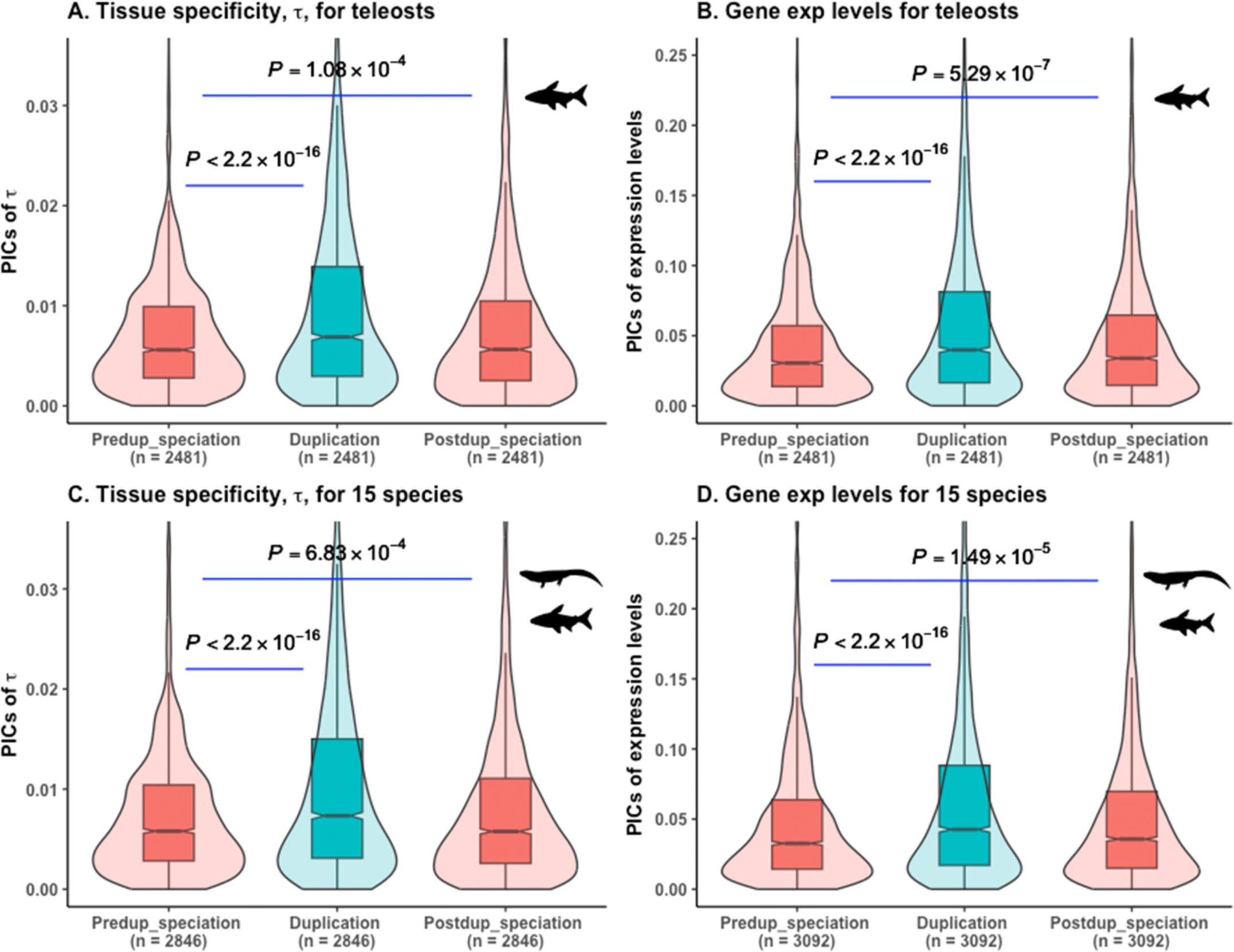
Modified phylogenetic approach to test the ortholog conjecture for teleosts and for vertebrates, respectively. We used the youngest duplication node along a path of a phylogeny for identification of pre-duplication and post-duplication speciations. For (A) and (B), we considered the contrasts of teleosts specific duplication and post-duplication speciation nodes in absence of teleost-specific pre-duplication speciation nodes contrasts. (C) and (D) are based on all 15 vertebrate species. PICs: Phylogenetic Independent Contrasts, exp: expression, Predup_speciation: Pre-duplication speciation, Postdup_speciation: Post-duplication speciation. Since contrasts are estimated for each tree, paired Wilcoxon test is used to compare the difference.

**Figure S9:**
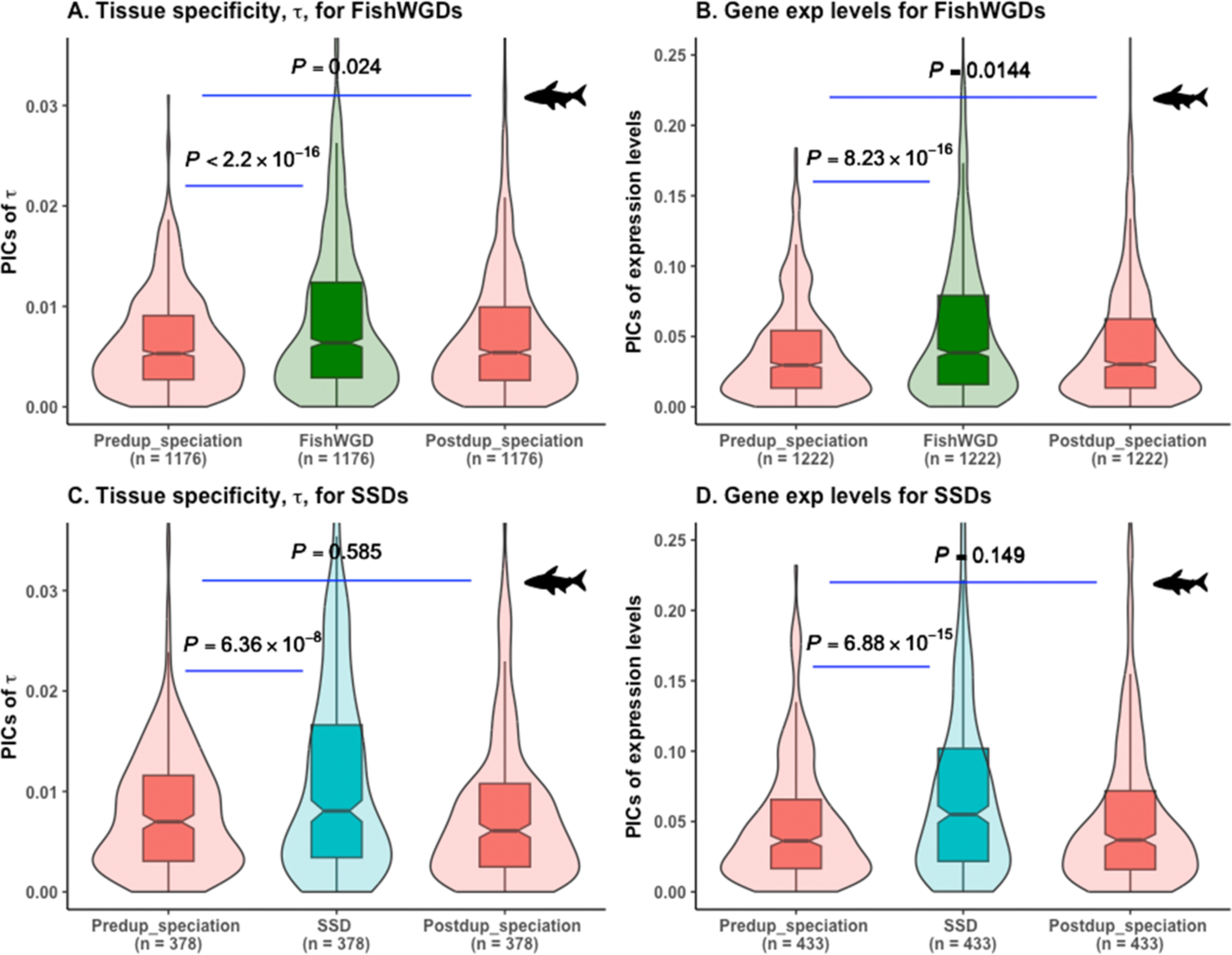
The ortholog conjecture tests on phylogenies with genome and gene duplicates in teleosts. 581, 610, 339 and 377 of trees passed our criteria of Fig. 4A in (A)-(D), respectively and were used for the analyses. We here used only teleost-specific duplication and post-duplication speciation nodes for contrasts analyses in absence of teleost-specific pre-duplication speciation nodes contrasts. We considered the youngest duplication node along a path of a phylogeny for identification of pre-duplication and post-duplication speciations. PICs: Phylogenetic independent contrasts, exp: expression, Predup_speciation: pre-duplication speciation, Postdup_speciation: post-duplication speciation, FishWGD: fish specific 3R whole genome duplications, 3RWGDs: ohnologs, SSDs: small-scale duplicates. Since contrasts are estimated for each tree, paired Wilcoxon test is used to compare the difference.

**Figure S10:**
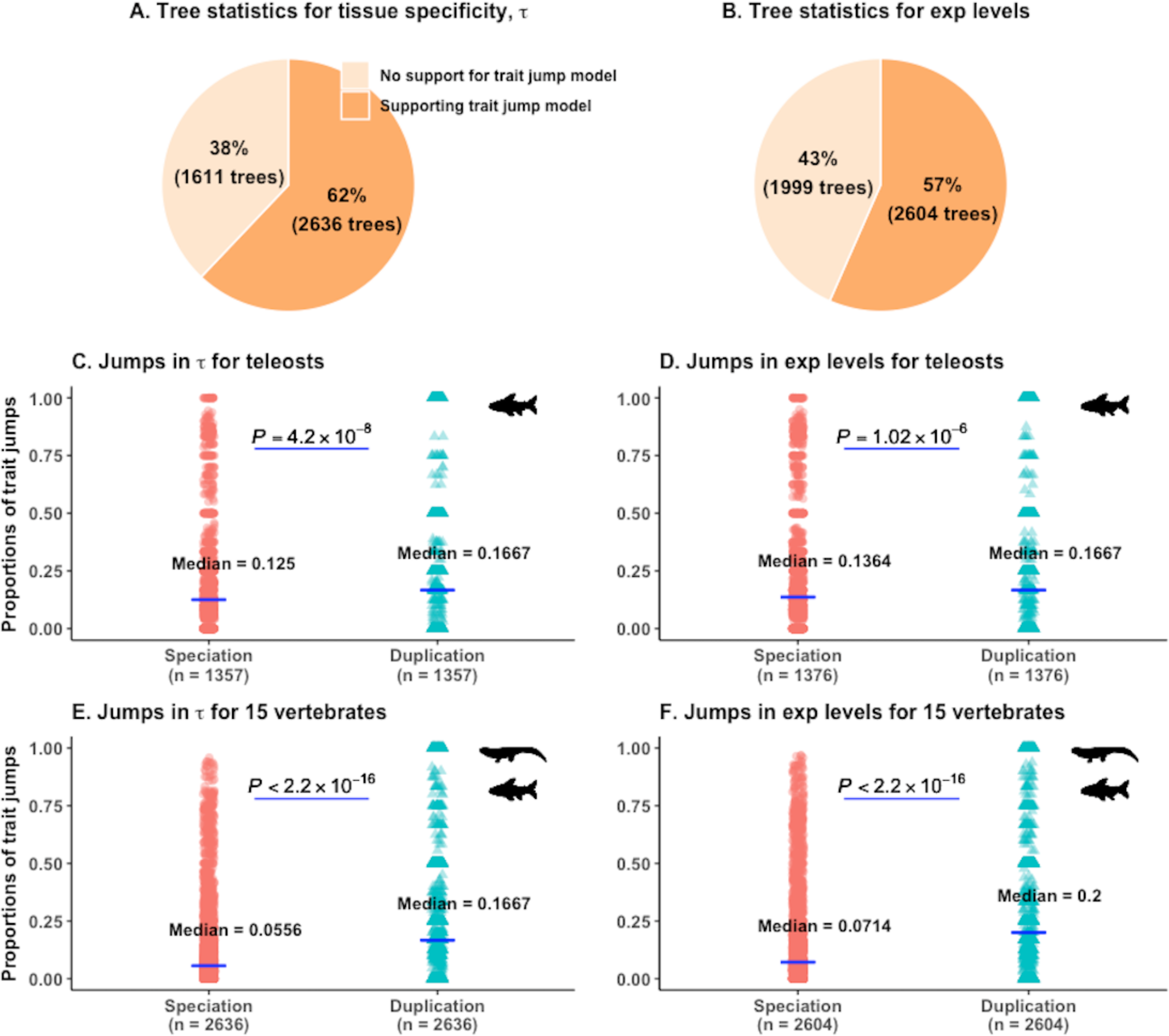
Analyses of trait jump models for τ and for average gene expression levels in 5 teleosts and in 15 vertebrates. We used a strict posterior probability cutoff of ≥ 0.7 as an evidence of trait jump. (A)-(B) Statistics of calibrated Brownian trees supporting jumps for τ and for average gene expression levels, respectively. Proportions of jumps in traits following speciations and duplication events are shown in (C)-(D) for teleosts and in (E)-(F) for vertebrates. 2636 trees supporting rapid jumps in τ and 2604 trees supporting jumps in average gene expression levels with the above-mentioned threshold were used for analyses. Since proportion of shifts in traits per branch of events is estimated for each tree, paired Wilcoxon test is used to compare the difference. ‘exp’: Average gene expression levels.

**Figure S11:**
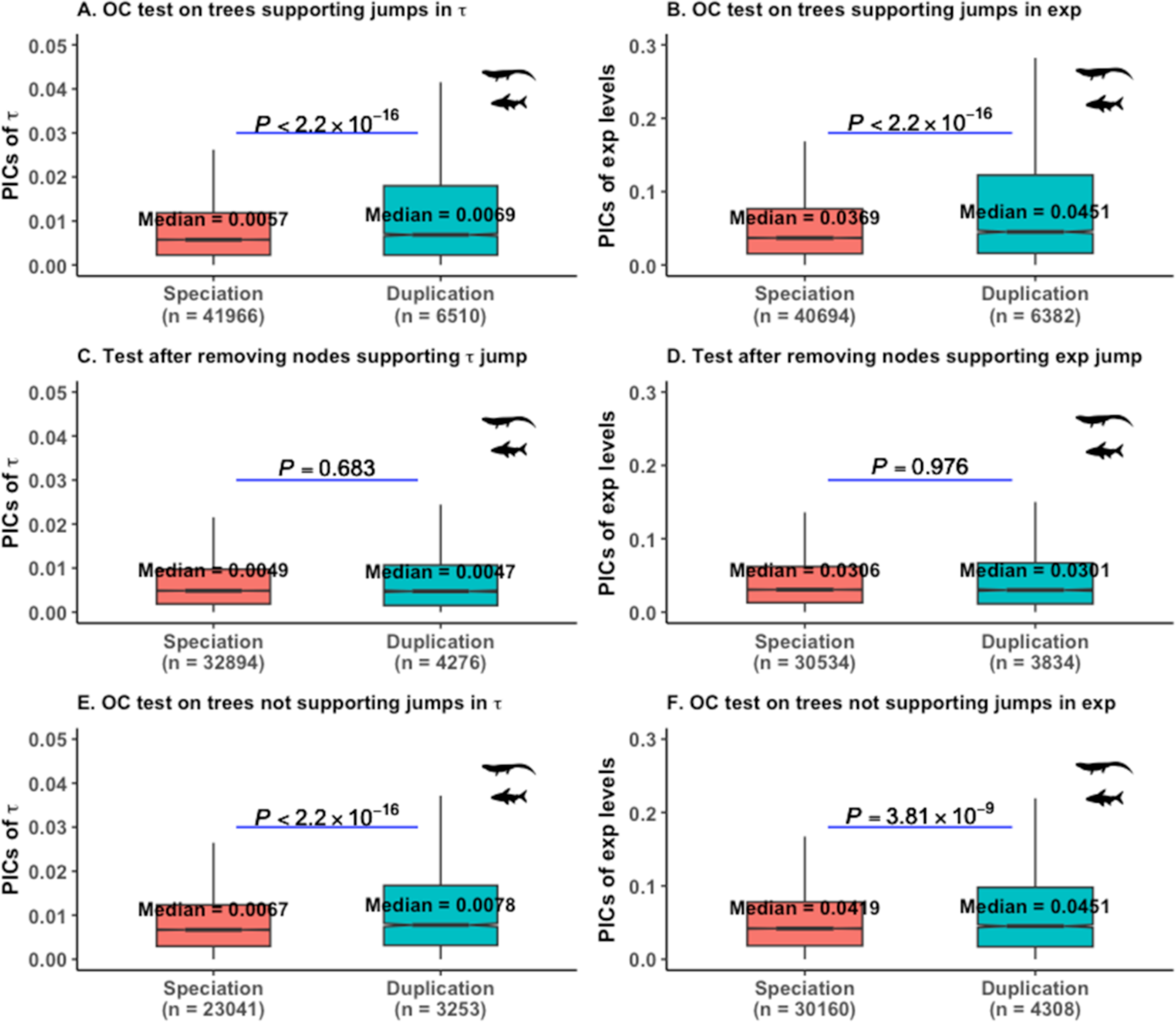
The ortholog conjecture (OC) tests on trees supporting rapid jump in τ and in average gene expression levels for 15 vertebrates. We used a strict branch jump posterior cutoff of ≥ 0.7 as an evidence of trait jump. Only teleosts-specific clades were used for the analyses. The ortholog conjecture tests on (A) 2636 calibrated contrast standardized trees supporting rapid jumps in τ, and (B) 2604 trees supporting jumps in average gene expression levels with the above-mentioned threshold along a phylogeny for our analyses. (C)-(D) The ortholog conjecture tests after removing contrasts of nodes whose daughter branch(es) experienced jumps in the corresponding trait for 2636 and 2604 trees, respectively. (E)-(F) The ortholog conjecture tests on 1611 and 1999 trees for which the trait jump model is not supported for τ and for average gene expression levels, respectively. *P* values are from Wilcoxon two-tailed tests.

**Figure S12:**
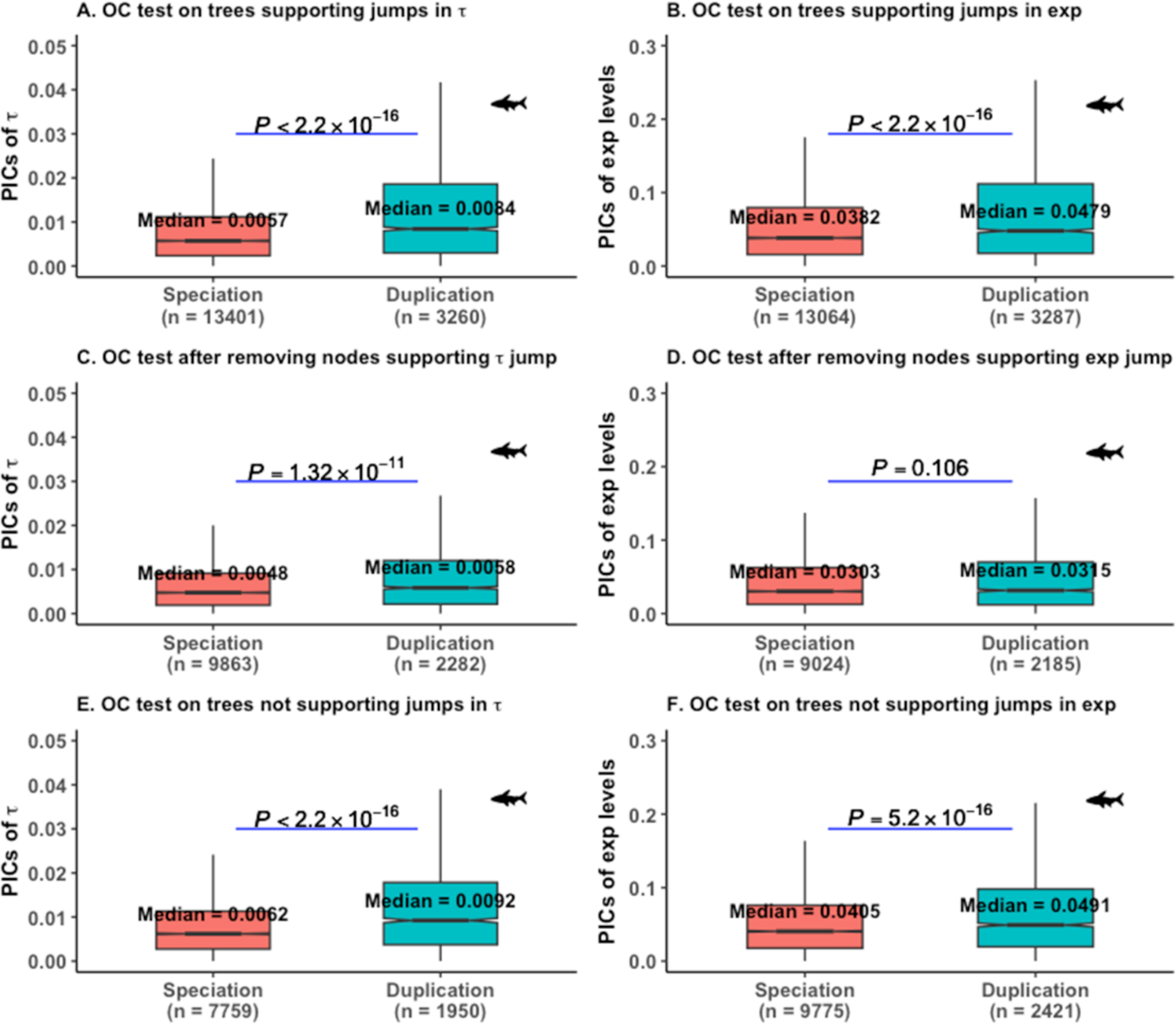
The ortholog conjecture (OC) tests on trees supporting rapid jump in τ and in average gene expression levels for 5 teleosts. We used a strict branch jump posterior cutoff of ≥ 0.7 as an evidence of trait jump. Only teleosts-specific clades were used for the analyses. (A) 2636 calibrated contrasts standardized trees supporting rapid jumps in τ, and (B) 2604 trees supporting jumps in average gene expression levels were used for the analyses. (C)-(D) The ortholog conjecture tests after removing contrasts of nodes whose daughter branch(es) experienced jumps in the corresponding trait for 2636 and 2604 trees, respectively. (E)-(F) The ortholog conjecture tests on 1611 and 1999 trees for which the trait jump model is not supported for τ and for average gene expression levels, respectively. *P* values are from Wilcoxon two-tailed tests.

**Figure S13:**
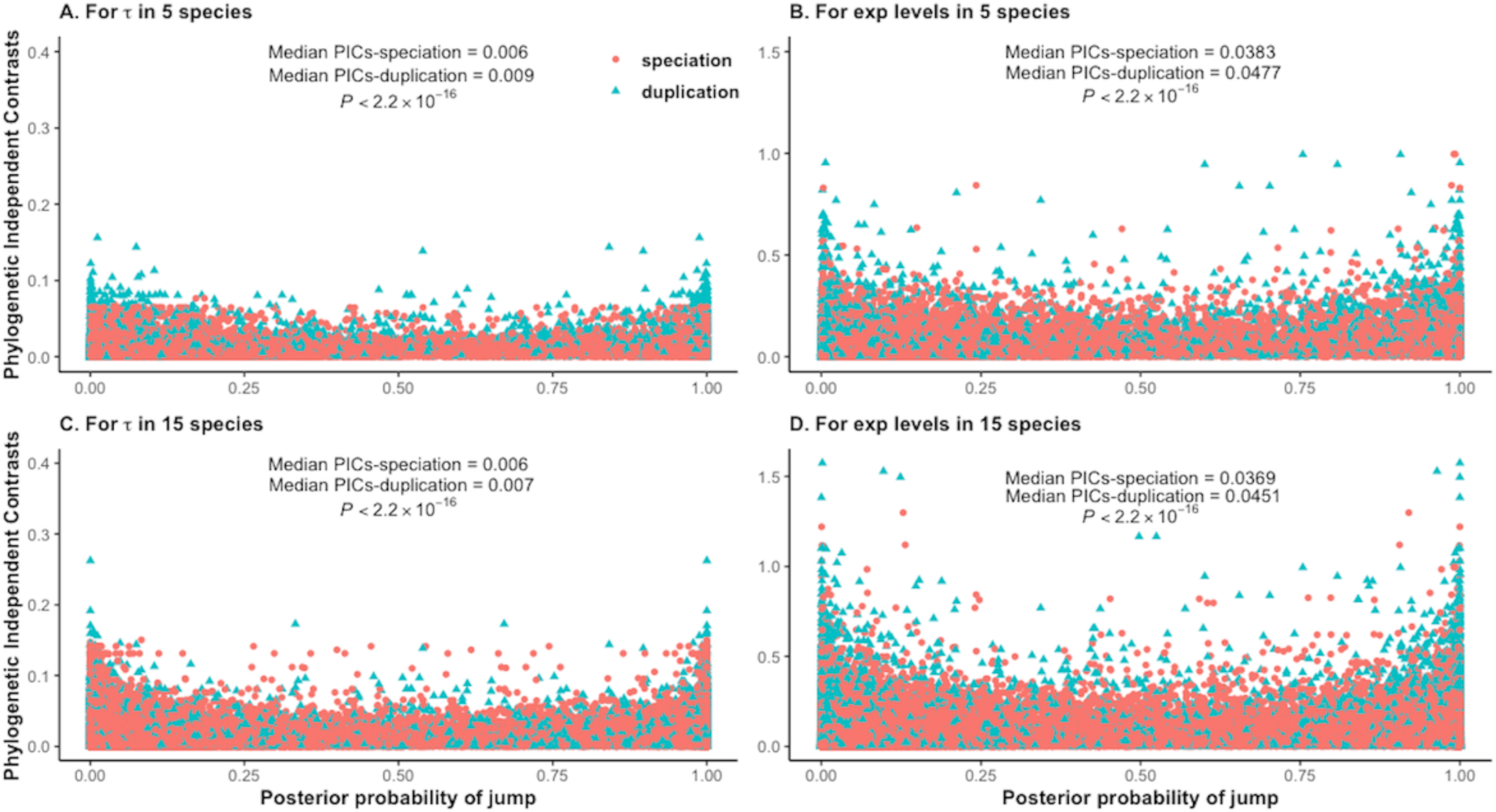
Distribution of the phylogenetic independent contrasts (PICs) for all the trees supporting rapid jump in τ and in average gene expression levels for teleosts and for vertebrates. For these analyses, we considered PICs of the nodes for a tree having a branch jump posterior probability > 0. Numbers of trees supporting jumps increased from 2636 to 2675 for τ and from 2604 to 2647 for average expression levels by altering the PP threshold from 0.7 to 0. *P* values are from Wilcoxon two-tailed tests.

**Figure S14:**
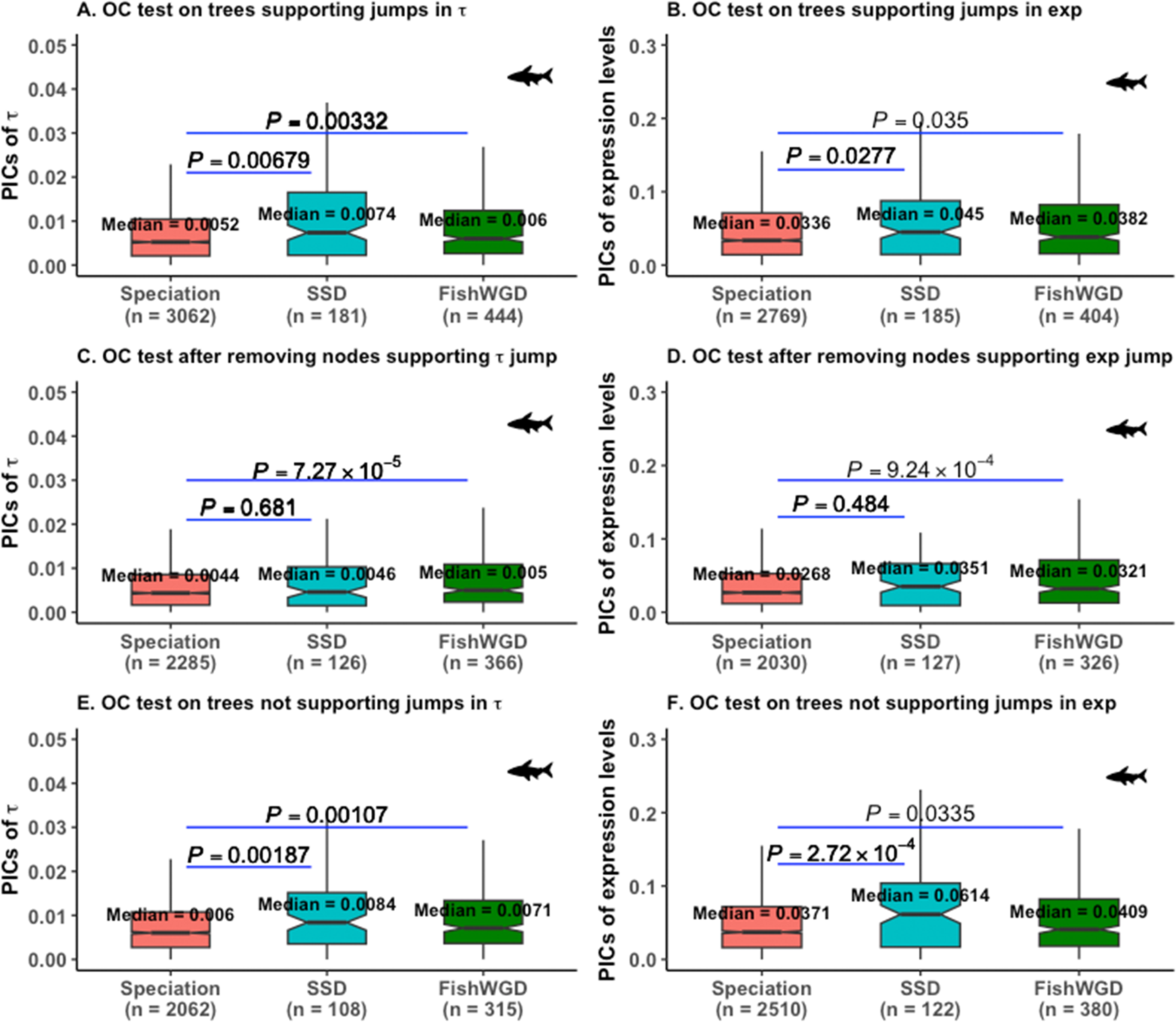
The ortholog conjecture (OC) tests on strict 3RWGD trees to analyze the jump results in τ and in average gene expression levels for teleosts. We used a strict branch jump posterior cutoff of ≥ 0.7 as an evidence of trait jump. Only teleosts-specific clades were used for the analyses. The ortholog conjecture tests on (A) 435 calibrated contrasts standardized strict 3RWGD trees supporting rapid jumps in τ, and on (B) 398 strict 3RWGD trees supporting jumps in average gene expression levels. (C)-(D) The ortholog conjecture tests after removing contrasts of nodes whose daughter branch(es) experienced jumps in the corresponding trait for 435 and 398 strict 3RWGD trees, respectively. (E)-(F) The ortholog conjecture tests on 311 and 376 strict 3RWGD trees for which the trait jump model is not supported for τ and for average gene expression levels, respectively. FishWGD: ohnologs, other duplicates refer to small scale duplicates. *P* values are from Wilcoxon two-tailed tests.

**Figure S15:**
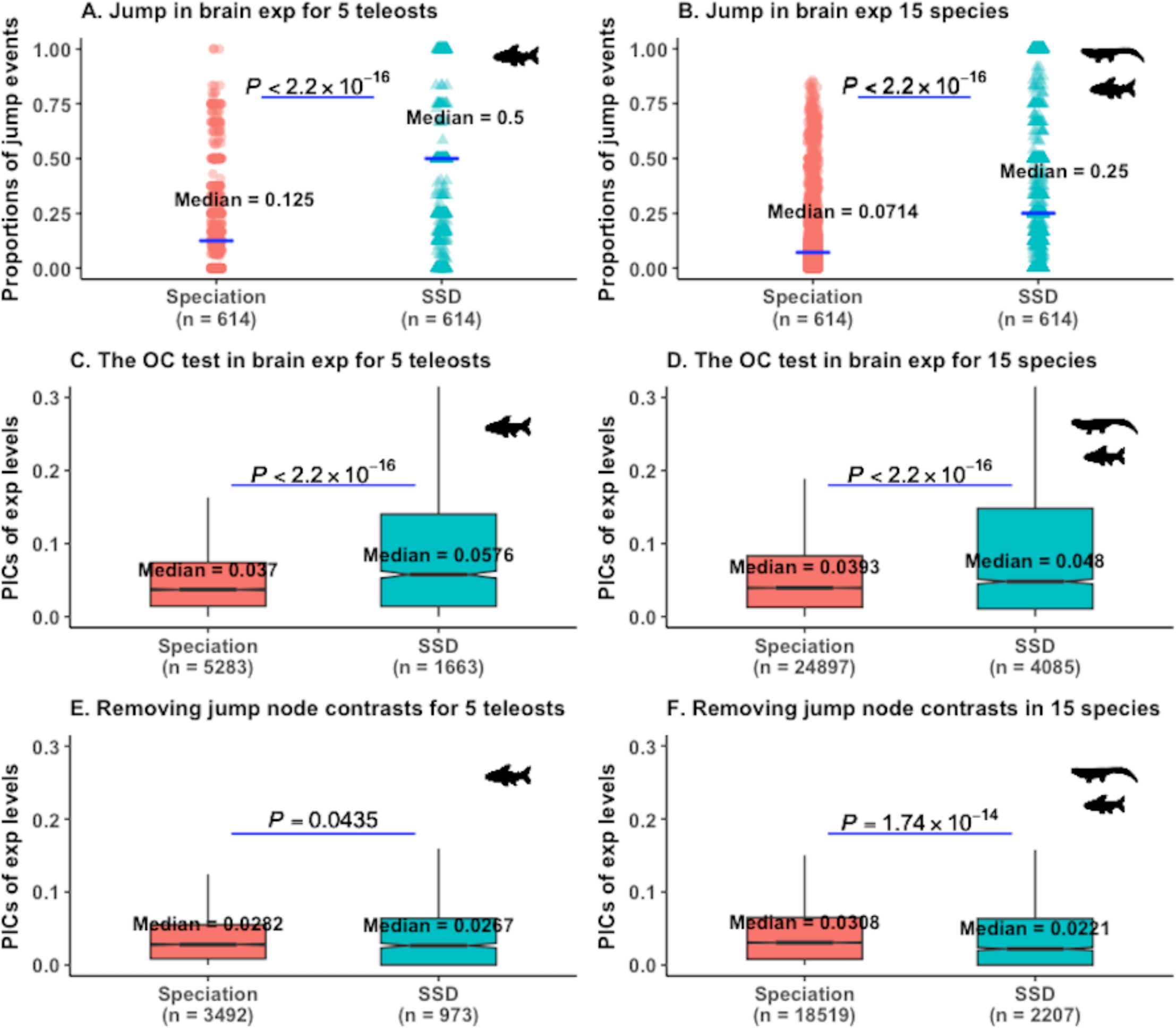
Analyses of rapid jumps in brain expression for strict SSD trees. We used a strict posterior probability cutoff of ≥ 0.7 as an evidence of trait jump. (A)-(B) Proportions of jumps in brain expressions following speciation and duplication events for teleosts and for vertebrates using 1752 trait jump supporting trees of Fig. 6A. Since proportion of shifts in traits per branch of events is estimated for each tree, paired Wilcoxon test is used to compare the difference for (A)-(B). (C)-(D) The ortholog conjecture (OC) tests on 1752 strict SSD trees to analyze the jump results in brain expressions for teleosts and for 15 vertebrates, respectively. (E)-(F) The ortholog conjecture tests after removing contrasts of nodes whose daughter branch(es) experienced jumps in the corresponding trait for 1752 strict SSD trees for 5 teleosts and for 15 vertebrates, respectively. *P* values are from Wilcoxon two-tailed tests for (C)-(F). ‘exp’: expression levels.

**Figure S16:**
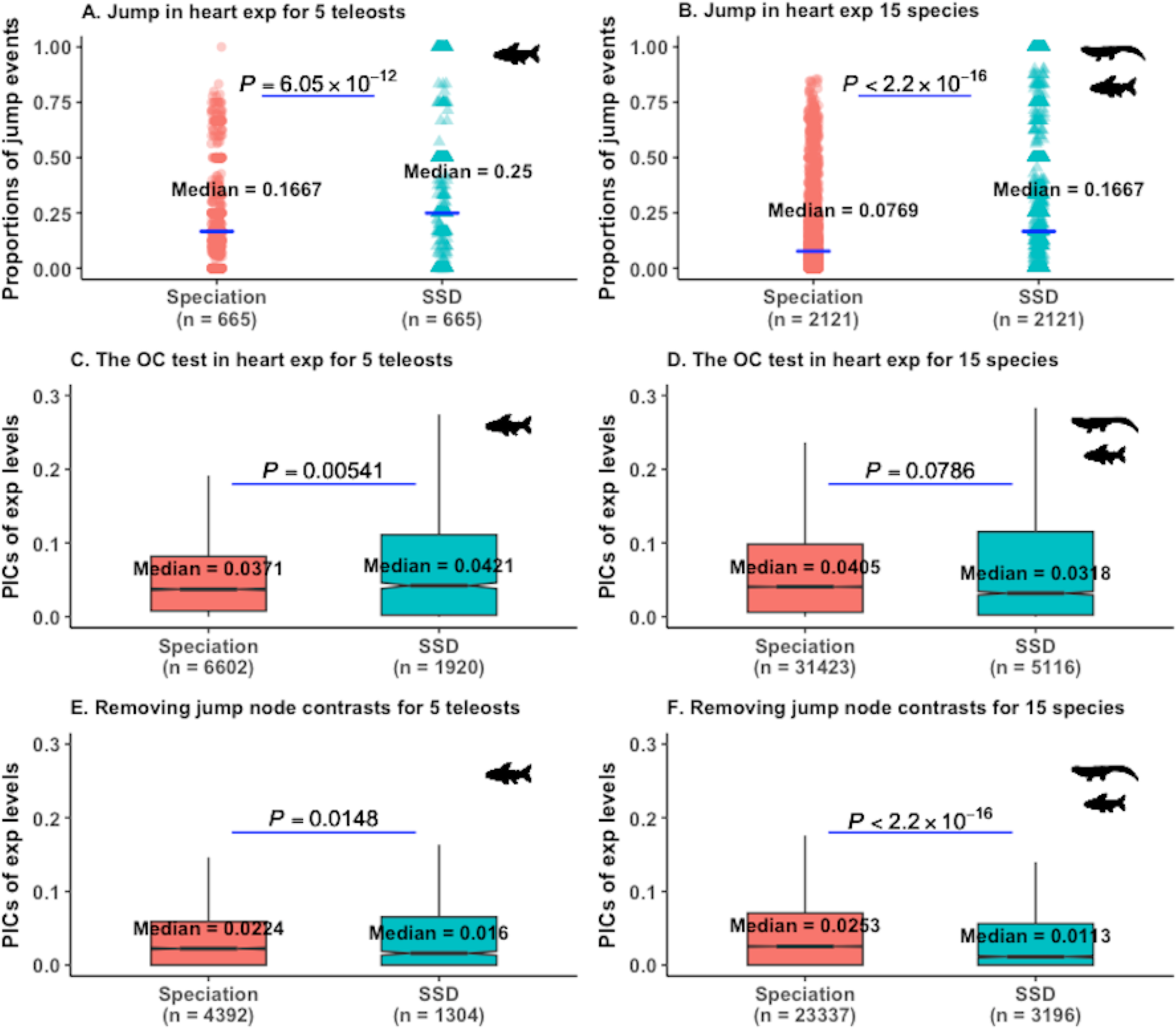
Analyses of rapid jumps in heart expression for strict SSD trees. We used a strict posterior probability cutoff of ≥ 0.7 as an evidence of trait jump. (A)-(B) Proportions of jumps in heart expressions following speciation and duplication events for teleosts and for vertebrates using 2135 trait jump supporting trees of Fig. 6B. Since proportion of shifts in traits per branch of events is estimated for each tree, paired Wilcoxon test is used to compare the difference for (A)-(B). (C)-(D) The ortholog conjecture (OC) tests on 2135 strict SSD trees to analyze the jump results in heart expressions for teleosts and for 15 vertebrates, respectively. (E)-(F) The ortholog conjecture tests after removing contrasts of nodes whose daughter branch(es) experienced jumps in the corresponding trait for 2135 strict SSD trees for 5 teleosts and for 15 vertebrates, respectively. *P* values are from Wilcoxon two-tailed tests for (C)-(F). ‘exp’: expression levels.

**Figure S17:**
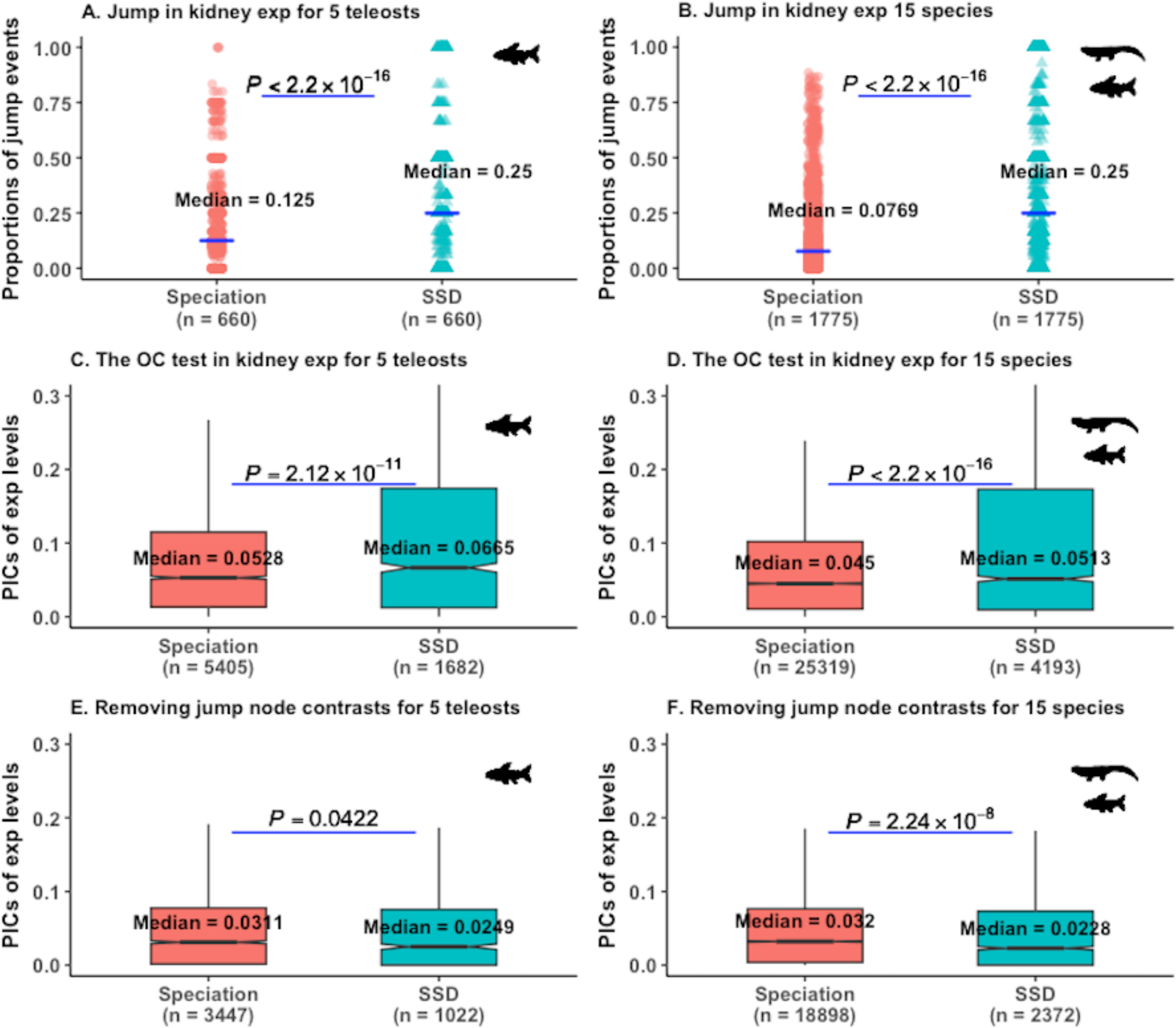
Analyses of rapid jumps in kidney expression for strict SSD trees. We used a strict posterior probability cutoff of ≥ 0.7 as an evidence of trait jump. (A)-(B) Proportions of jumps in kidney expressions following speciation and duplication events for teleosts and for vertebrates using 1788 trait jump supporting trees of Fig. 6C. Since proportion of shifts in traits per branch of events is estimated for each tree, paired Wilcoxon test is used to compare the difference for (A)-(B). (C)-(D) The ortholog conjecture (OC) tests on 1788 strict SSD trees to analyze the jump results in kidney expressions for teleosts and for 15 vertebrates, respectively. (E)-(F) The ortholog conjecture tests after removing contrasts of nodes whose daughter branch(es) experienced jumps in the corresponding trait for 1788 strict SSD trees for 5 teleosts and for 15 vertebrates, respectively. *P* values are from Wilcoxon two-tailed tests for (C)-(F). ‘exp’: expression levels.

**Figure S18:**
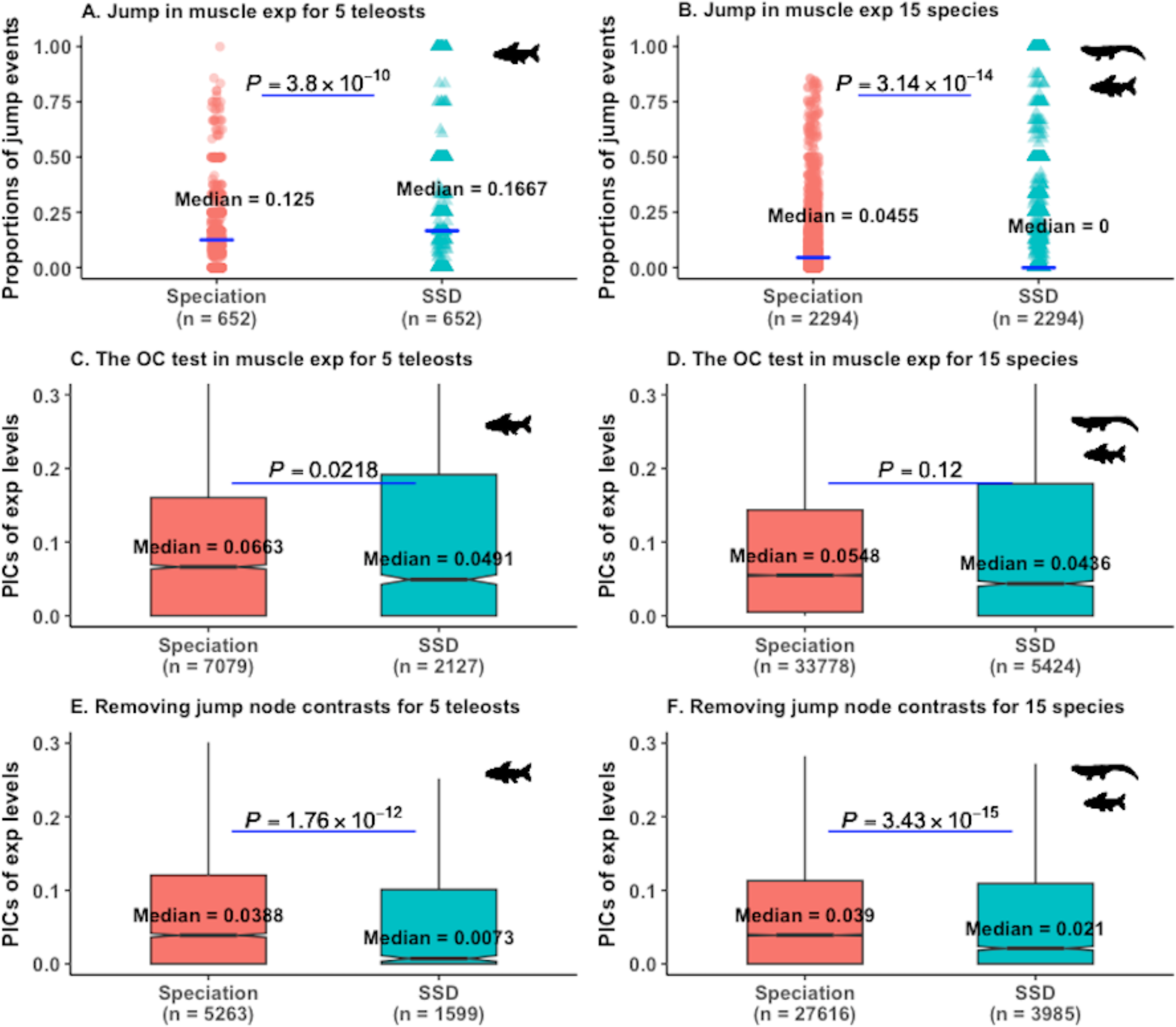
Analyses of rapid jumps in muscle expression for strict SSD trees. We used a strict posterior probability cutoff of ≥ 0.7 as an evidence of trait jump. (A)-(B) Proportions of jumps in muscle expressions following speciation and duplication events for teleosts and for vertebrates using 2319 trait jump supporting trees of Fig. 6D. Since proportion of shifts in traits per branch of events is estimated for each tree, paired Wilcoxon test is used to compare the difference for (A)-(B). (C)-(D) The ortholog conjecture (OC) tests on 2319 strict SSD trees to analyze the jump results in muscle expressions for teleosts and for 15 vertebrates, respectively. (E)-(F) The ortholog conjecture tests after removing contrasts of nodes whose daughter branch(es) experienced jumps in the corresponding trait for 2319 strict SSD trees for 5 teleosts and for 15 vertebrates, respectively. *P* values are from Wilcoxon two-tailed tests for (C)-(F). ‘exp’: expression levels.

**Figure S19:**
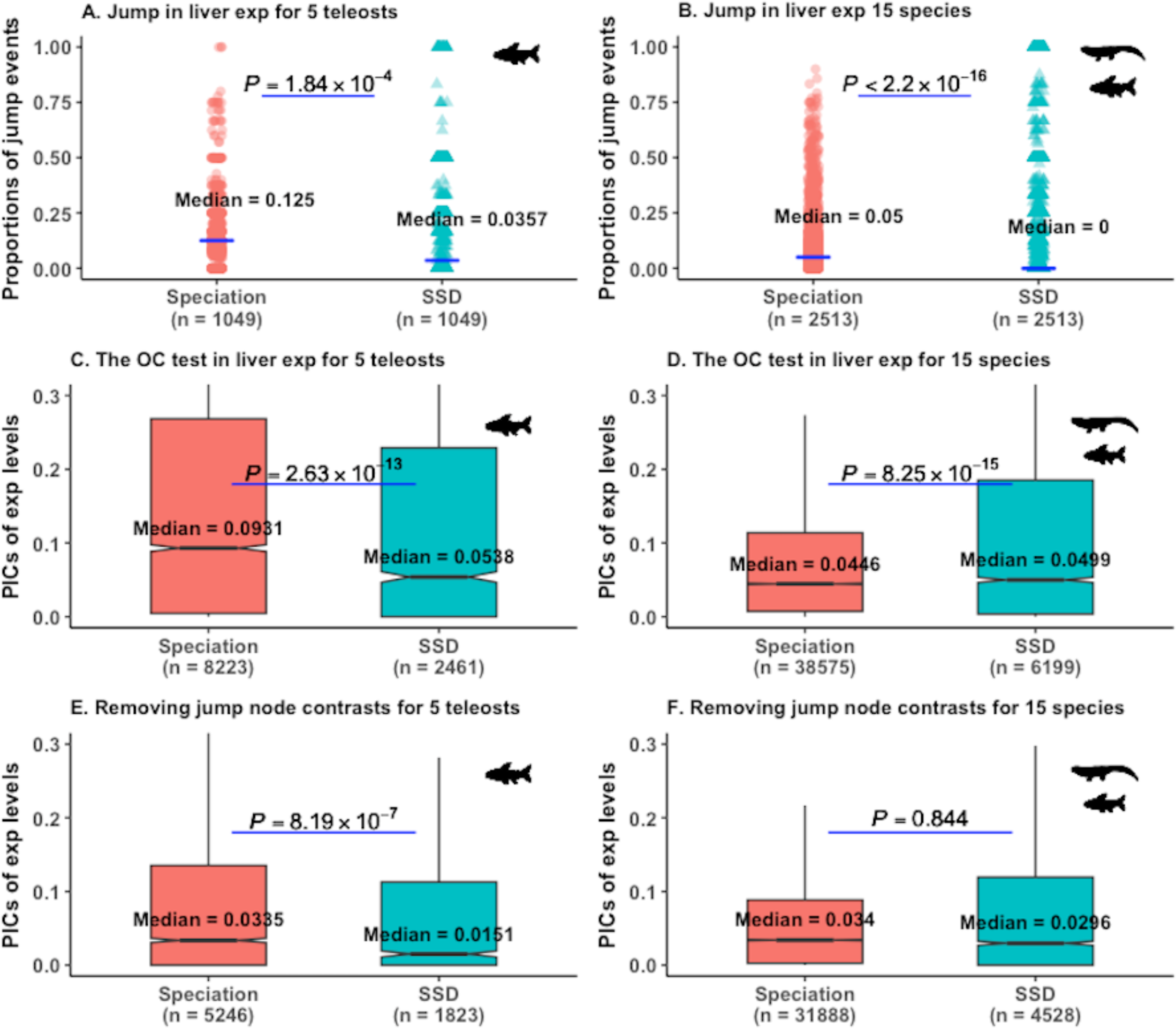
Analyses of rapid jumps in liver expression for strict SSD trees. We used a strict posterior probability cutoff of ≥ 0.7 as an evidence of trait jump. (A)-(B) Proportions of jumps in liver expressions following speciation and duplication events for teleosts and for vertebrates using 2641 trait jump supporting trees of Fig. 6E. Since proportion of shifts in traits per branch of events is estimated for each tree, paired Wilcoxon test is used to compare the difference for (A)-(B). (C)-(D) The ortholog conjecture (OC) tests on 2641 strict SSD trees to analyze the jump results in liver expressions for teleosts and for 15 vertebrates, respectively. (E)-(F) The ortholog conjecture tests after removing contrasts of nodes whose daughter branch(es) experienced jumps in the corresponding trait for 2641 strict SSD trees for 5 teleosts and for 15 vertebrates, respectively. *P* values are from Wilcoxon two-tailed tests for (C)-(F). ‘exp’: expression levels.

**Figure S20:**
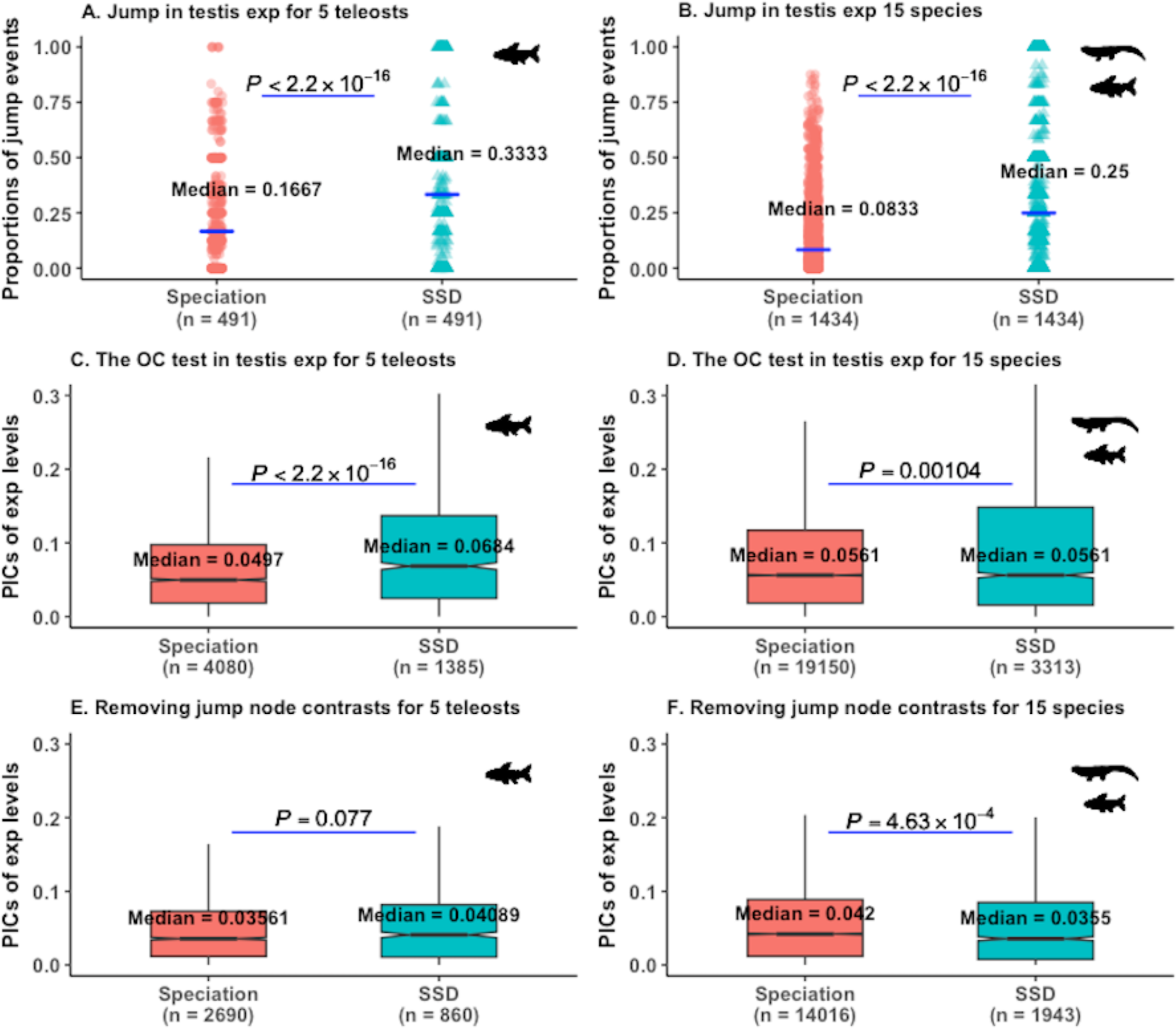
Analyses of rapid jumps in testis expression for strict SSD trees. We used a strict posterior probability cutoff of ≥ 0.7 as an evidence of trait jump. (A)-(B) Proportions of jumps in testis expressions following speciation and duplication events for teleosts and for vertebrates using 1441 trait jump supporting trees of Fig. 6F. Since proportion of shifts in traits per branch of events is estimated for each tree, paired Wilcoxon test is used to compare the difference for (A)-(B). (C)-(D) The ortholog conjecture (OC) tests on 1441 strict SSD trees to analyse the jump results in testis expressions for teleosts and for 15 vertebrates, respectively. (E)-(F) The ortholog conjecture tests after removing contrasts of nodes whose daughter branch(es) experienced jumps in the corresponding trait for 1441 strict SSD trees for 5 teleosts and for 15 vertebrates, respectively. *P* values are from Wilcoxon two-tailed tests for (C)-(F). ‘exp’: expression levels.

**Figure S21:**
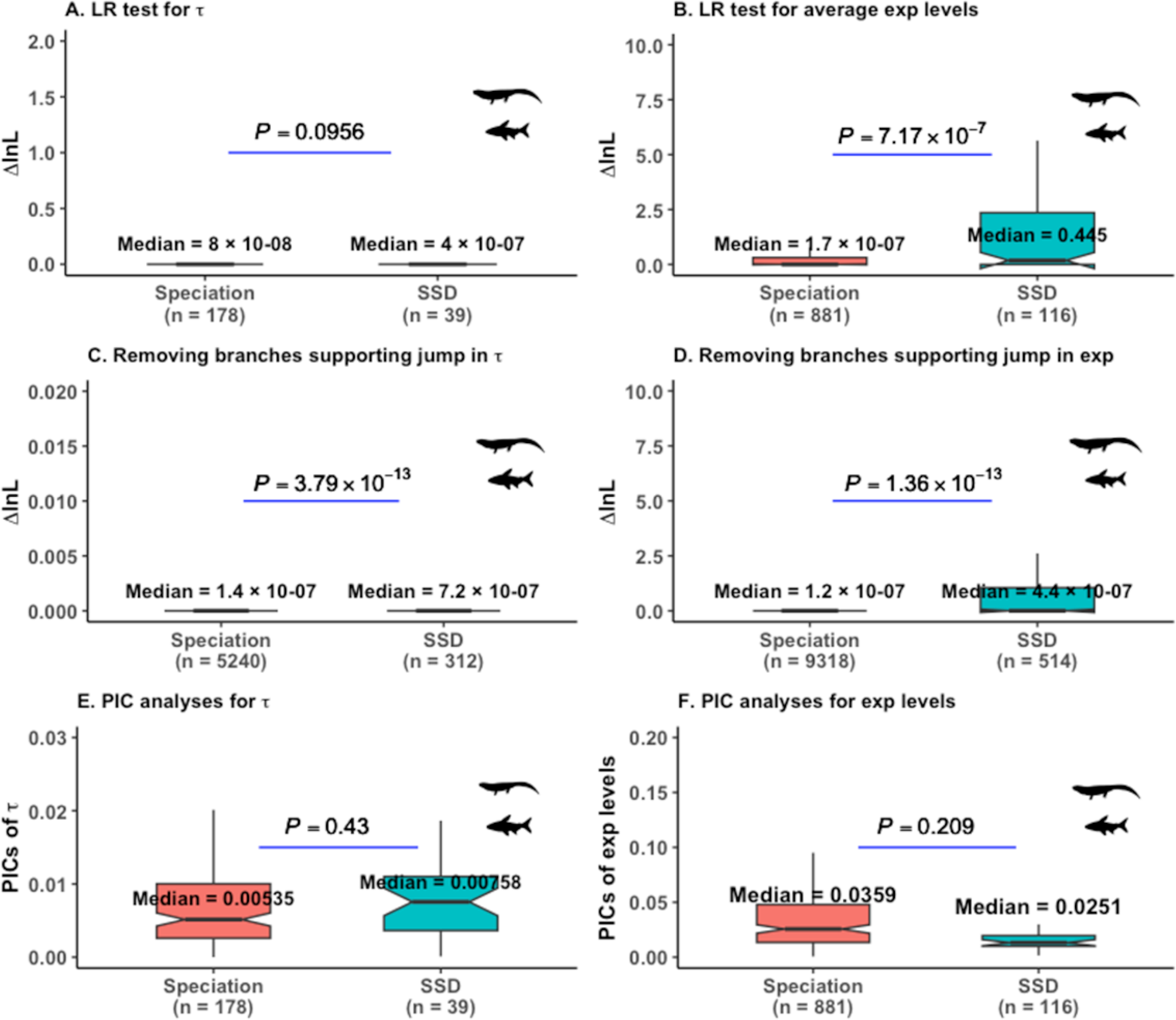
Positive selection and trait divergence analyses for contrast standardized strict SSD trees in all 15 vertebrates. We only used phylogenies supporting an evolutionary trait jump model for these analyses. We used a strict posterior probability cutoff of ≥ 0.7 to identify branches with rapid jumps in traits. exp: Gene expression, PICs: phylogenetic independent contrasts, LRT: likelihood ratio test.

**Table S1:** Statistics on 1575 and 1619 strict SSD trees by using posterior probability (PP) threshold of 0.5 for. τ **and for average expression levels, respectively.**

| Events | | Number of<br>branches with<br>evolutionary<br>trait jumps (PP<br>$\geq 0.5$) | Number of<br>branches with no<br>inferred jump<br>(PP $< 0.5$) | Proportions<br>of trait<br>jumps | Two-sample test for<br>equality of<br>proportions, $\chi^2$ test<br>with Yates correction |
| --- | --- | --- | --- | --- | --- |
| For $\tau$ in 5<br>teleost-specific clades | **Speciation** | 865 | 3807 | 0.185 | $\chi^2 = 74.21$ on $df = 1$,<br>$P < 1 \times 10^{-4}$ |
|  | **SSD** | 581 | 1505 | 0.278 |  |
| For average<br>expression<br>levels in 5<br>teleost-specific clades | **Speciation** | 1307 | 4177 | 0.238 | $\chi^2 = 113.24$ on $df = 1$,<br>$P < 1 \times 10^{-4}$ |
|  | **SSD** | 900 | 1654 | 0.352 |  |
| For $\tau$ in all 15<br>vertebrate<br>clades | **Speciation** | 8194 | 40152 | 0.169 | $\chi^2 = 630.51$ on $df = 1$,<br>$P < 1 \times 10^{-4}$ |
|  | **SSD** | 2149 | 5219 | 0.292 |  |
| For average<br>expression<br>levels in all 15<br>vertebrate<br>clades | **Speciation** | 5245 | 22841 | 0.187 | $\chi^2 = 597.56$ on $df = 1$,<br>$P < 1 \times 10^{-4}$ |
|  | **SSD** | 1562 | 2947 | 0.346 |  |

*Note:* Numbers of strict SSD trees with jumps increased from 1557 to 1575 and from 1593 to 1619 trees by altering the PP threshold from 0.7 to 0.5 for $\tau$ and for average expression levels, respectively.

**Table S2:**
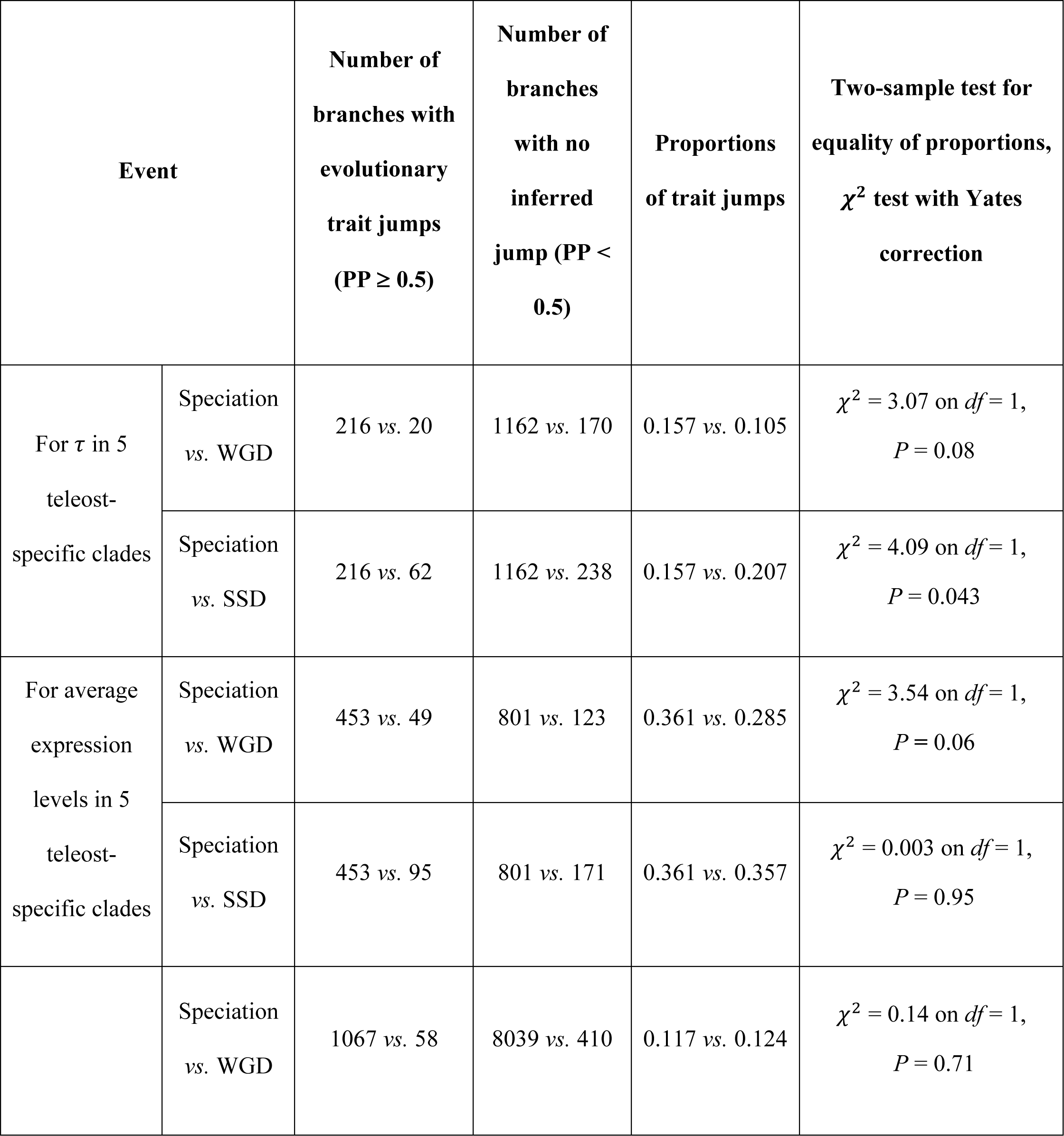
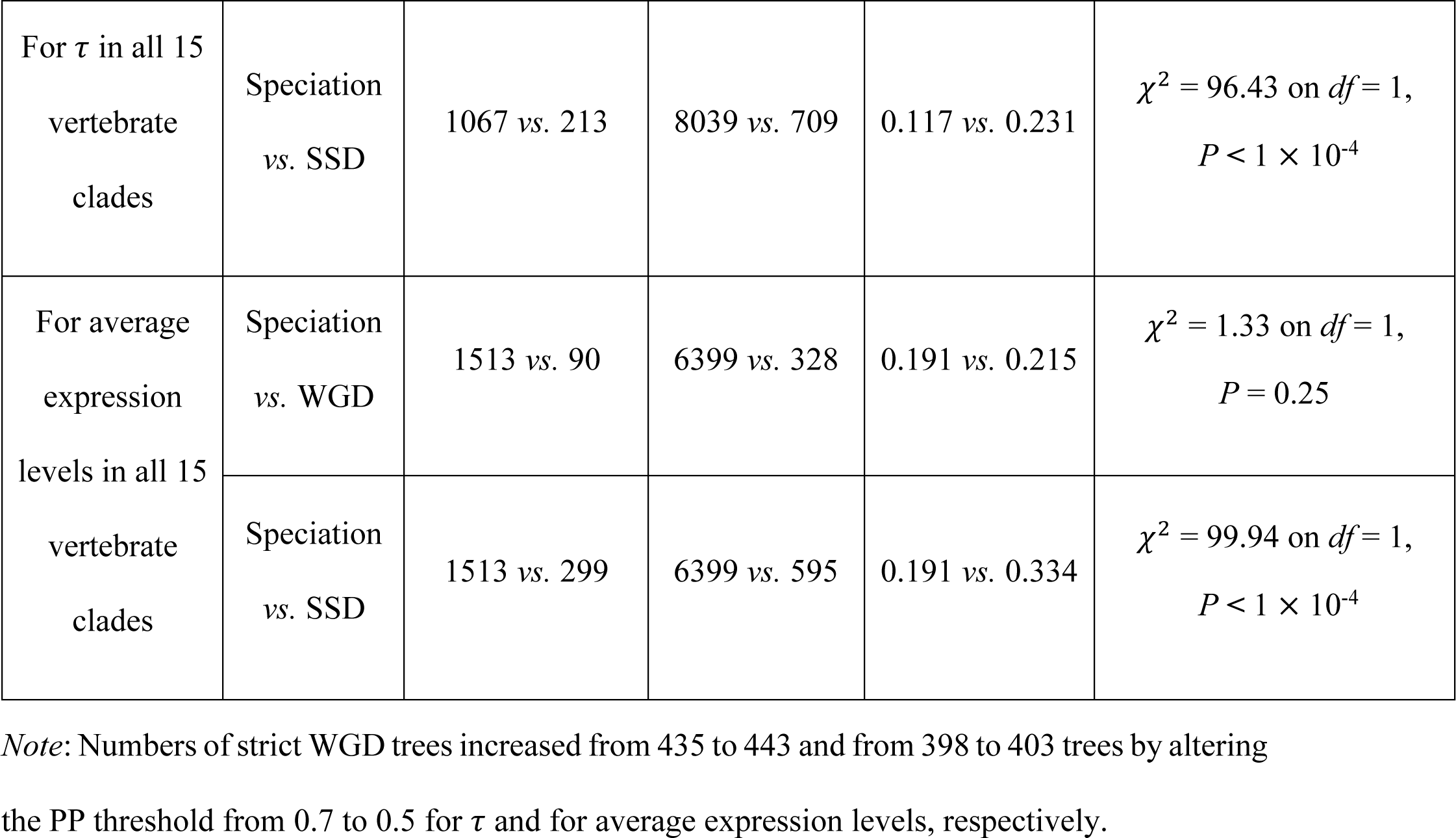
Statistics on 443 and 403 strict WGD trees by using posterior probability (PP) threshold of 0.5 for τ and for average expression levels, respectively.

## Notes

https://github.com/tbegum/phylogenetic-modeling-of-evolutionary_trait_jumps

https://zenodo.org/records/10418895

### Competing Interest Statement

The authors have declared no competing interest.

### Summary of Updates

Both the use of phylogenetic independent contrasts and of the jump model were clarified, and controls were added.

## References

Altenhoff AM, Studer RA, Robinson-Rechavi M, Dessimoz C. 2012. Resolving the ortholog conjecture: Orthologs tend to be weakly, but significantly, more similar in function than paralogs. PLoS Comput Biol 8:e1002514. Available from: https://www.ncbi.nlm.nih.gov/pubmed/22615551

Andrew S. 2019.FastQC: A quality control tool for high throughput sequence data 2010.

Anisimova M, Yang Z. 2007. Multiple hypothesis testing to detect lineages under positive selection that affects only a few sites. Mol Biol Evol 24:1219–1228. Available from: https://www.ncbi.nlm.nih.gov/pubmed/17339634

Antoine Lucas DE with contributions by, Tuszynski J, Bengtsson H, Urbanek S, Frasca M, Lewis B, Stokely M, Muehleisen H, Murdoch D, Hester J, et al. 2018. Digest: Create compact hash digests of r objects. Available from: https://CRAN.R-project.org/package=digest

Assis R, Bachtrog D. 2013. Neofunctionalization of young duplicate genes in drosophila. Proc Natl Acad Sci U S A 110:17409–17414. Available from: https://www.ncbi.nlm.nih.gov/pubmed/24101476

Assis R, Bachtrog D. 2015. Rapid divergence and diversification of mammalian duplicate gene functions. BMC Evol Biol 15:138. Available from: https://www.ncbi.nlm.nih.gov/pubmed/26173681

Auguie B. 2017. GridExtra: Miscellaneous functions for “grid” graphics. Available from: https://CRAN.R-project.org/package=gridExtra

Bastian FB, Roux J, Niknejad A, Comte A, Fonseca Costa SS, Farias TM de, Moretti S, Parmentier G, Laval VR de, Rosikiewicz M, et al. 2021. The Bgee suite: Integrated curated expression atlas and comparative transcriptomics in animals. Nucleic Acids Res 49:D831–D847. Available from: https://www.ncbi.nlm.nih.gov/pubmed/33037820

Beaulieu JM, Jhwueng DC, Boettiger C, O’Meara BC. 2012. Modeling stabilizing selection: Expanding the ornstein-uhlenbeck model of adaptive evolution. Evolution 66:2369–2383. Available from: https://www.ncbi.nlm.nih.gov/pubmed/22834738

Begum T, Robinson-Rechavi M. 2021. Special care is needed in applying phylogenetic comparative methods to gene trees with speciation and duplication nodes. Mol Biol Evol 38:1614– 1626. Available from: https://www.ncbi.nlm.nih.gov/pubmed/33169790

Begum T, Serrano-Serrano ML, Robinson-Rechavi M. 2021. Performance of a phylogenetic independent contrast method and an improved pairwise comparison under different scenarios of trait evolution after speciation and duplication. Methods Ecol Evol 12:1875–1887.

Berthelot C, Brunet F, Chalopin D, Juanchich A, Bernard M, Noël B, Bento P, Da Silva C, Labadie K, Alberti A, et al. 2014. The rainbow trout genome provides novel insights into evolution after whole-genome duplication in vertebrates. Nat Commun 5:3657. Available from: https://www.ncbi.nlm.nih.gov/pubmed/24755649

Blomberg SP, Garland T, Ives AR. 2003. Testing for phylogenetic signal in comparative data: Behavioral traits are more labile. Evolution 57:717–745. Available from: https://www.ncbi.nlm.nih.gov/pubmed/12778543

Bokma F. 2008. Detection of “punctuated equilibrium” by bayesian estimation of speciation and extinction rates, ancestral character states, and rates of anagenetic and cladogenetic evolution on a molecular phylogeny. Evolution 62:2718–2726. Available from: https://www.ncbi.nlm.nih.gov/pubmed/18752617

Bolger AM, Lohse M, Usadel B. 2014. Trimmomatic: A flexible trimmer for illumina sequence data. Bioinformatics 30:2114–2120. Available from: https://www.ncbi.nlm.nih.gov/pubmed/24695404

Braasch I, Gehrke AR, Smith JJ, Kawasaki K, Manousaki T, Pasquier J, Amores A, Desvignes T, Batzel P, Catchen J, et al. 2016. The spotted gar genome illuminates vertebrate evolution and facilitates human-teleost comparisons. Nat Genet 48:427–437. Available from: https://www.ncbi.nlm.nih.gov/pubmed/26950095

Brawand D, Soumillon M, Necsulea A, Julien P, Csárdi G, Harrigan P, Weier M, Liechti A, Aximu-Petri A, Kircher M, et al. 2011. The evolution of gene expression levels in mammalian organs. Nature 478:343–348. Available from: https://www.ncbi.nlm.nih.gov/pubmed/22012392

Bray NL, Pimentel H, Melsted P, Pachter L. 2016. Near-optimal probabilistic rna-seq quantification. Nat Biotechnol 34:525–527.

Brunet FG, Roest Crollius H, Paris M, Aury JM, Gibert P, Jaillon O, Laudet V, Robinson-Rechavi M. 2006. Gene loss and evolutionary rates following whole-genome duplication in teleost fishes. Mol Biol Evol 23:1808–1816. Available from: https://www.ncbi.nlm.nih.gov/pubmed/16809621

Chamberlain S. 2018. Rphylopic: Get’silhouettes’ of’organisms’ from’Phylopic’. R package version 0.2. 0.

Chen J, Swofford R, Johnson J, Cummings BB, Rogel N, Lindblad-Toh K, Haerty W, Palma FD, Regev A. 2019. A quantitative framework for characterizing the evolutionary history of mammalian gene expression. Genome Res 29:53–63. Available from: https://www.ncbi.nlm.nih.gov/pubmed/30552105

Chen X, Zhang J. 2012. The ortholog conjecture is untestable by the current gene ontology but is supported by rna sequencing data. PLoS Comput Biol 8:e1002784. Available from: https://www.ncbi.nlm.nih.gov/pubmed/23209392

Conant GC, Wagner A. 2003. Asymmetric sequence divergence of duplicate genes. Genome Res 13:2052–2058. Available from: https://www.ncbi.nlm.nih.gov/pubmed/12952876

Cooper N, Thomas GH, FitzJohn RG. 2016. Shedding light on the ‘dark side’ of phylogenetic comparative methods. Methods Ecol Evol 7:693–699. Available from: https://www.ncbi.nlm.nih.gov/pubmed/27499839

Cusack BP, Wolfe KH. 2007. Not born equal: Increased rate asymmetry in relocated and retrotransposed rodent gene duplicates. Mol Biol Evol 24:679–686. Available from: https://www.ncbi.nlm.nih.gov/pubmed/17179139

Daub JT, Moretti S, Davydov II, Excoffier L, Robinson-Rechavi M. 2017. Detection of pathways affected by positive selection in primate lineages ancestral to humans. Mol Biol Evol 34:391– 1402. Available from: 10.1093/molbev/msx083

Davesne D, Friedman M, Schmitt AD, Fernandez V, Carnevale G, Ahlberg PE, Sanchez S, Benson RBJ. 2021. Fossilized cell structures identify an ancient origin for the teleost whole-genome duplication. Proc Natl Acad Sci U S A 118. Available from: https://www.ncbi.nlm.nih.gov/pubmed/34301898

David KT, Oaks JR, Halanych KM. 2020. Patterns of gene evolution following duplications and speciations in vertebrates. PeerJ 8:e8813. Available from: https://www.ncbi.nlm.nih.gov/pubmed/32266119

Dennis MY, Nuttle X, Sudmant PH, Antonacci F, Graves TA, Nefedov M, Rosenfeld JA, Sajjadian S, Malig M, Kotkiewicz H, et al. 2012. Evolution of human-specific neural srgap2 genes by incomplete segmental duplication. Cell 149:912–922. Available from: https://www.ncbi.nlm.nih.gov/pubmed/22559943

Diaz-Uriarte R, Garland T. 1996. Testing hypotheses of correlated evolution using phylogenetically independent contrasts: Sensitivity to deviations from brownian motion. Syst Biol 45:27–47.

Díaz-Uriarte R, Garland T. 1998. Effects of branch length errors on the performance of phylogenetically independent contrasts. Syst Biol 47:654–672. Available from: https://www.ncbi.nlm.nih.gov/pubmed/12066309

Dougherty ML, Nuttle X, Penn O, Nelson BJ, Huddleston J, Baker C, Harshman L, Duyzend MH, Ventura M, Antonacci F, et al. 2017. The birth of a human-specific neural gene by incomplete duplication and gene fusion. Genome Biol 18:49. Available from: https://www.ncbi.nlm.nih.gov/pubmed/28279197

Duchen P, Alfaro ML, Rolland J, Salamin N, Silvestro D. 2021. On the effect of asymmetrical trait inheritance on models of trait evolution. Syst Biol 70:376–388. Available from: https://www.ncbi.nlm.nih.gov/pubmed/32681798

Duchen P, Leuenberger C, Szilágyi SM, Harmon L, Eastman J, Schweizer M, Wegmann D. 2017. Inference of evolutionary jumps in large phylogenies using lévy processes. Syst Biol 66:950–963. Available from: https://www.ncbi.nlm.nih.gov/pubmed/28204787

Dunn CW, Zapata F, Munro C, Siebert S, Hejnol A. 2018. Pairwise comparisons across species are problematic when analyzing functional genomic data. Proc Natl Acad Sci U S A 115:E409– E417. Available from: https://www.ncbi.nlm.nih.gov/pubmed/29301966

Eastman JM, Alfaro ME, Joyce P, Hipp AL, Harmon LJ. 2011. A novel comparative method for identifying shifts in the rate of character evolution on trees. Evolution 65:3578–3589. Available from: https://www.ncbi.nlm.nih.gov/pubmed/22133227

Felsenstein J. 1985. Confidence limits on phylogenies: An approach using the bootstrap. Evolution 39:783–791. Available from: https://www.ncbi.nlm.nih.gov/pubmed/28561359

Freckleton R, Harvey P, Pagel M. 2002. Phylogenetic analysis and comparative data: A test and review of evidence. Am Nat 160:712–726.

Freckleton RP, Harvey PH. 2006. Detecting non-brownian trait evolution in adaptive radiations. PLoS Biol 4:e373. Available from: https://www.ncbi.nlm.nih.gov/pubmed/17090217

Fukushima K, Pollock DD. 2020. Amalgamated cross-species transcriptomes reveal organ-specific propensity in gene expression evolution. Nat Commun 11:4459. Available from: https://www.ncbi.nlm.nih.gov/pubmed/32900997

Gabaldón T, Koonin EV. 2013. Functional and evolutionary implications of gene orthology. Nat Rev Genet 14:360–366. Available from: https://www.ncbi.nlm.nih.gov/pubmed/23552219

Gao Y, Wu M. 2022. Microbial genomic trait evolution is dominated by frequent and rare pulsed evolution. Sci Adv 8:eabn1916. Available from: DOI: 10.1126/sciadv.abn19

Garland T. 1992. Rate tests for phenotypic evolution using phylogenetically independent contrasts. Am Nat 140:509–519. Available from: https://www.ncbi.nlm.nih.gov/pubmed/19426053

Gillard, G.B., Grønvold, L., Røsæg, L.L. et al. 2021. Comparative regulomics supports pervasive selection on gene dosage following whole genome duplication. Genome Biol 22:1–18.

Gout JF, Lynch M. 2015. Maintenance and loss of duplicated genes by dosage subfunctionalization. Mol Biol Evol 32:2141–2148. Available from: https://www.ncbi.nlm.nih.gov/pubmed/25908670

Grafen A. 1989. The phylogenetic regression. Philos Trans R Soc Lond B Biol Sci 326:119–157. Available from: https://www.ncbi.nlm.nih.gov/pubmed/2575770

Gu X, Su Z. 2007. Tissue-driven hypothesis of genomic evolution and sequence-expression correlations. Proc Natl Acad Sci U S A 104:2779–2784. Available from: https://www.ncbi.nlm.nih.gov/pubmed/17301236

Gu X, Zhang Z, Huang W. 2005. Rapid evolution of expression and regulatory divergences after yeast gene duplication. Proc Natl Acad Sci U S A 102:707–712. Available from: https://www.ncbi.nlm.nih.gov/pubmed/15647348

Guangchuang Y. 2018. Treeio: Base classes and functions for phylogenetic tree input and output. Available from: https://guangchuangyu.github.io/software/treeio

Guangchuang Y, David S, Huachen Z, Yi G, Tommy T-YL. 2017. Ggtree: An r package for visualization and annotation of phylogenetic trees with their covariates and other associated data. Methods Ecol Evol 8:28–36.

Guschanski K, Warnefors M, Kaessmann H. 2017. The evolution of duplicate gene expression in mammalian organs. Genome Res 27:1461–1474. Available from: https://www.ncbi.nlm.nih.gov/pubmed/28743766

Han MV, Demuth JP, McGrath CL, Casola C, Hahn MW. 2009. Adaptive evolution of young gene duplicates in mammals. Genome Res 19:859–867. Available from: https://www.ncbi.nlm.nih.gov/pubmed/19411603

Hansen TF. 1997. Stabilizing selection and the comparative analysis of adaptation. Evolution 51:1341–1351. Available from: https://www.ncbi.nlm.nih.gov/pubmed/28568616

Hedges SB, Dudley J, Kumar S. 2006. TimeTree: A public knowledge-base of divergence times among organisms. Bioinformatics 22:2971–2972. Available from: https://www.ncbi.nlm.nih.gov/pubmed/17021158

Holland PW, Marlétaz F, Maeso I, Dunwell TL, Paps J. 2017. New genes from old: Asymmetric divergence of gene duplicates and the evolution of development. Philos Trans R Soc Lond B Biol Sci 372. Available from: https://www.ncbi.nlm.nih.gov/pubmed/27994121

Kaessmann, H, Vinckenbosch N, Long M. 2009. RNA-based gene duplication: mechanistic and evolutionary insights. Nat Rev Genet 10:19–31. Available from: 10.1038/nrg2487

Khabbazian M, Kriebel R, Rohe K, Ané C. 2016. Fast and accurate detection of evolutionary shifts in ornstein-uhlenbeck models. Methods Ecol Evol 7:811–824.

Kim SH, Yi SV. 2006. Correlated asymmetry of sequence and functional divergence between duplicate proteins of saccharomyces cerevisiae. Mol Biol Evol 23:1068–1075. Available from: https://www.ncbi.nlm.nih.gov/pubmed/16510556

Koonin EV. 2005. Orthologs, paralogs, and evolutionary genomics. Annu Rev Genet 39:309–338. Available from: https://www.ncbi.nlm.nih.gov/pubmed/16285863

Kryuchkova-Mostacci N, Robinson-Rechavi M. 2016. Tissue-specificity of gene expression diverges slowly between orthologs, and rapidly between paralogs. PLoS Comput Biol 12:e1005274. Available from: https://www.ncbi.nlm.nih.gov/pubmed/28030541

Kryuchkova-Mostacci N, Robinson-Rechavi M. 2017. A benchmark of gene expression tissue-specificity metrics. Brief Bioinform 18:205–214. Available from: 10.1093/bib/bbw008

Landis MJ, Schraiber JG. 2017. Pulsed evolution shaped modern vertebrate body sizes. Proc Natl Acad Sci USA 114: 13224–13229. Available from: https://www.ncbi.nlm.nih.gov/pubmed/29114046

Landis MJ, Schraiber JG, Liang M. 2013. Phylogenetic analysis using lévy processes: Finding jumps in the evolution of continuous traits. Syst Biol 62:193–204. Available from: https://www.ncbi.nlm.nih.gov/pubmed/23034385

Lemmon AR, Moriarty EC. 2004. The importance of proper model assumption in Bayesian phylogenetics. Syst Biol 1:265–77.

Lien S, Koop BF, Sandve SR, Miller JR, Kent MP, Nome T, Hvidsten TR, Leong JS, Minkley DR, Zimin A, et al. 2016. The atlantic salmon genome provides insights into rediploidization. Nature 533:200–205. Available from: https://www.ncbi.nlm.nih.gov/pubmed/27088604

Lynch M, Katju V. 2004. The altered evolutionary trajectories of gene duplicates. Trends Genet 20:544–549. Available from: https://www.ncbi.nlm.nih.gov/pubmed/15475113

Martins E, Hansen T. 1997. Phylogenies and the comparative method: A general approach to incorporating phylogenetic information into the analysis of interspecific data. Am Nat 149:646– 667.

Martin FJ, Amode MR, Aneja A, Austine-Orimoloye O, Azov AG, et al. 2023. Ensembl 2023. Nucleic Acids Res 51: D933–D941. Available from 10.1093/nar/gkac958

Moretti S, Laurenczy B, Gharib WH, Castella B, Kuzniar A, Schabauer H, Studer RA, Valle M, Salamin N, Stockinger H, Robinson-Rechavi M. 2014. Selectome update: quality control and computational improvements to a database of positive selection, Nucleic Acids Res 42:D917– D921. 10.1093/nar/gkt1065

Nehrt NL, Clark WT, Radivojac P, Hahn MW. 2011. Testing the ortholog conjecture with comparative functional genomic data from mammals. PLoS Comput Biol 7:e1002073. Available from: https://www.ncbi.nlm.nih.gov/pubmed/21695233

Nguyen NTT, Vincens P, Dufayard JF, Roest Crollius H, Louis A. 2022. Genomicus in 2022: Comparative tools for thousands of genomes and reconstructed ancestors. Nucleic Acids Res 50:D1025–D1031. Available from: https://www.ncbi.nlm.nih.gov/pubmed/34792170

Ohno S. 1970. Evolution by gene duplication. New York (EUA). Springer-Verlag.

O’Meara BC, Ané C, Sanderson MJ, Wainwright PC. 2006. Testing for different rates of continuous trait evolution using likelihood. Evolution 60:922–933. Available from: https://www.ncbi.nlm.nih.gov/pubmed/16817533

Orme D. 2018. The caper package: Comparative analysis of phylogenetics and evolution in R. Available from: https://cran.r-project.org/web/packages/caper/vignettes/caper.pdf

Pagel M. 1999. Inferring the historical patterns of biological evolution. Nature 401:877–884.

Panchin AY, Gelfand MS, Ramensky VE, Artamonova II. 2010. Asymmetric and non-uniform evolution of recently duplicated human genes. Biol Direct 5:54. Available from: https://www.ncbi.nlm.nih.gov/pubmed/20825637

Paradis E, Claude J, Strimmer K. 2004. APE: Analyses of phylogenetics and evolution in R language. Bioinformatics 20:289–290. Available from: https://www.ncbi.nlm.nih.gov/pubmed/14734327

Parey E, Louis A, Cabau C, Guiguen Y, Roest Crollius H, Berthelot C. 2020. Synteny-guided resolution of gene trees clarifies the functional impact of whole-genome duplications. Mol Biol Evol 37:3324–3337. Available from: https://www.ncbi.nlm.nih.gov/pubmed/32556216

Parey E, Louis A, Montfort J, Guiguen Y, Roest Crollius H, Berthelot C. 2022. An atlas of fish genome evolution reveals delayed rediploidization following the teleost whole-genome duplication. Genome Res. Available from: https://www.ncbi.nlm.nih.gov/pubmed/35961774

Parey, E, Alexandra L, Jérôme M, et al. 2023. Genome structures resolve the early diversification of teleost fishes. Science 379:572–3575. Available from: https://www.ncbi.nlm.nih.gov/pubmed/36758078

Pasquier J, Braasch I, Batzel P, Cabau C, Montfort J, Nguyen T, Jouanno E, Berthelot C, Klopp C, Journot L, et al. 2017. Evolution of gene expression after whole-genome duplication: New insights from the spotted gar genome. J Exp Zool B Mol Dev Evol 328:709–721. Available from: https://www.ncbi.nlm.nih.gov/pubmed/28944589

Pasquier J, Cabau C, Nguyen T, Jouanno E, Severac D, Braasch I, Journot L, Pontarotti P, Klopp C, Postlethwait JH, et al. 2016. Gene evolution and gene expression after whole genome duplication in fish: The PhyloFish database. BMC Genomics 17:368. Available from: https://www.ncbi.nlm.nih.gov/pubmed/27189481

Pegueroles C, Laurie S, Albà MM. 2013. Accelerated evolution after gene duplication: A time-dependent process affecting just one copy. Mol Biol Evol 30:1830–1842. Available from: https://www.ncbi.nlm.nih.gov/pubmed/23625888

Pennell MW, Eastman JM, Slater GJ, Brown JW, Uyeda JC, FitzJohn RG, Alfaro ME, Harmon LJ. 2014. Geiger v2.0: An expanded suite of methods for fitting macroevolutionary models to phylogenetic trees. Bioinformatics 30:2216–2218. Available from: https://www.ncbi.nlm.nih.gov/pubmed/24728855

Pich I Roselló O, Kondrashov FA. 2014. Long-term asymmetrical acceleration of protein evolution after gene duplication. Genome Biol Evol 6:1949–1955. Available from: https://www.ncbi.nlm.nih.gov/pubmed/25070510

Pilmann Kotěrová A, Santos F, Bejdová Š. et al. 2024. Prioritizing a high posterior probability threshold leading to low error rate over high classification accuracy: the validity of MorphoPASSE software for cranial morphological sex estimation in a contemporary population. Int J Legal Med. Available from: 10.1007/s00414-024-03215-1

Purvis A, Rambaut A. 1995. Comparative analysis by independent contrasts (caic): An apple macintosh application for analysing comparative data. Comput Appl Biosci 11:247–251. Available from: https://www.ncbi.nlm.nih.gov/pubmed/7583692

R Core Team. 2018. R: A language and environment for statistical computing. Vienna, Austria: R Foundation for Statistical Computing. Available from: https://www.R-project.org/

Revell LJ. 2012. Phytools: An r package for phylogenetic comparative biology (and other things). Methods Ecol Evol 3:217–223.

Revell LJ. 2014. Ancestral character estimation under the threshold model from quantitative genetics, Evolution 68:743–759. Available from: 10.1111/evo.12300

Rogozin IB, Managadze D, Shabalina SA, Koonin EV. 2014. Gene family level comparative analysis of gene expression in mammals validates the ortholog conjecture. Genome Biol Evol 6:754–762. Available from: https://www.ncbi.nlm.nih.gov/pubmed/24610837

Rohlfs RV, Nielsen R. 2015. Phylogenetic ANOVA: The expression variance and evolution model for quantitative trait evolution. Syst Biol 64:695–708. Available from: https://www.ncbi.nlm.nih.gov/pubmed/26169525

Romero IG, Ruvinsky I, Gilad Y. 2012. Comparative studies of gene expression and the evolution of gene regulation. Nat Rev Genet 13:505–516. Available from: https://www.ncbi.nlm.nih.gov/pubmed/22705669

Sandve SR, Rohlfs RV, Hvidsten TR. 2018. Subfunctionalization versus neofunctionalization after whole-genome duplication. Nat Genet 50:908–909. Available from: https://www.ncbi.nlm.nih.gov/pubmed/29955176

Scannell DR, Wolfe KH. 2008. A burst of protein sequence evolution and a prolonged period of asymmetric evolution follow gene duplication in yeast. Genome Res 18:137–147. Available from: https://www.ncbi.nlm.nih.gov/pubmed/18025270

Simpson G. 1944. Tempo and mode in evolution. A wartime book. New York: Columbia University Press.

Singh PP, Isambert H. 2020. OHNOLOGS v2: A comprehensive resource for the genes retained from whole genome duplication in vertebrates. Nucleic Acids Res 48:D724–D730. Available from: https://www.ncbi.nlm.nih.gov/pubmed/31612943

Stamboulian M, Guerrero RF, Hahn MW, Radivojac P. 2020. The ortholog conjecture revisited: The value of orthologs and paralogs in function prediction. Bioinformatics 36:i219–i226. Available from: https://www.ncbi.nlm.nih.gov/pubmed/32657391

Storey JD, Tibshirani R. 2003. Statistical significance for genomewide studies. Proc Natl Acad Sci U S A 100:9440–9445. Available from: https://www.ncbi.nlm.nih.gov/pubmed/12883005

Studer RA, Robinson-Rechavi M. 2009. How confident can we be that orthologs are similar, but paralogs differ? Trends Genet 25:210–216. Available from: https://www.ncbi.nlm.nih.gov/pubmed/19368988

Urbanek S. 2013. Png: Read and write png images. Available from: https://CRAN.R-project.org/package=png

Uyeda J, Hansen T, Arnold S, Pienaar J. 2011. The million-year wait for macroevolutionary bursts. Proc. Natl Acad. Sci. USA 108:15908–15913.

Uyeda JC, Harmon LJ. 2014. A novel bayesian method for inferring and interpreting the dynamics of adaptive landscapes from phylogenetic comparative data. Syst Biol 63:902–918. Available from: https://www.ncbi.nlm.nih.gov/pubmed/25077513

Warnes GR, Bolker B, Lumley T. 2018. Gtools: Various r programming tools. Available from: https://CRAN.R-project.org/package=gtools

Wickham H. 2016. Ggplot2: Elegant graphics for data analysis. Springer-Verlag New York Available from: https://ggplot2.tidyverse.org

Wickham H. 2017. Tidyverse: Easily install and load the ‘tidyverse’. Available from: https://CRAN.R-project.org/package=tidyverse

Wickham H. 2019. Stringr: Simple, consistent wrappers for common string operations. Available from: https://CRAN.R-project.org/package=stringr

Wickham H, Francois R, Henry L, Muller K. 2017. Dplyr: A grammar of data manipulation. Available from: https://CRAN.R-project.org/package=dplyr

Yanai I, Benjamin H, Shmoish M, Chalifa-Caspi V, Shklar M, Ophir R, Bar-Even A, Horn-Saban S, Safran M, Domany E, et al. 2005. Genome-wide midrange transcription profiles reveal expression level relationships in human tissue specification. Bioinformatics 21:650–659. Available from: <Go to ISI>://WOS:000227241200012

